# Personalized multi-assay profiling of respiratory motile ciliopathies and mRNA therapy

**DOI:** 10.64898/2026.05.21.726963

**Authors:** Georgia-Nefeli Ithakisiou, Pim Cleijpool, Henriette H.M. Dreyer, Bram M. Bosch, Wessel Hornman, Daniëlle Hoenselaar, Marinos Tziouvelis, Aranka Gerritsen, Matthew B. Smith, Loes A. den Hertog-Oosterhoff, Rumpa B. Bhattacharjee, Zechen Wang, T. Noelle Lombana, Brandon A. Wustman, Cornelis K. van der Ent, Karin M. de Winter-Groot, Sam F.B. van Beuningen, Eric G. Haarman, Tamara Paff, Jeffrey M. Beekman, Gimano D. Amatngalim, Bahar Yetkin-Arik

## Abstract

**Introduction:** Impaired motile cilia function contributes to many respiratory disorders, but therapies targeting this cellular defect are currently lacking. Personalized airway epithelial models combined with quantitative, complementary ciliary assays can pave the way for the development of such therapies. However, existing airway epithelial cultures often show variable ciliogenesis, and ciliary function is frequently assessed using a single assay that does not capture the phenotypic heterogeneity of ciliary dysfunction. Here, we established a personalized, multi-assay *in vitro* platform using human nasal epithelial cells (HNECs) to assess ciliary function and therapeutic response, using primary ciliary dyskinesia (PCD) as a model disease.

**Methods:** HNECs from 8 healthy individuals and 13 individuals with PCD carrying distinct disease-associated variants were obtained by nasal brushing. Cells were differentiated under optimized conditions, including γ-secretase/Notch and BMP pathway inhibitors and a low liquid-liquid interface, to generate highly ciliated 2D epithelial cultures. Ciliary function was assessed using ciliary beat frequency, bead transport, and apical-out nasal organoid rotation assays. Therapeutic rescue was assessed in HNECs harboring *DNAI1* alterations using *DNAI1* mRNA-loaded lipid nanoparticles.

**Results:** Optimized differentiation yielded reproducibly multiciliated HNEC cultures. The multi-assay platform distinguished healthy from PCD-derived HNECs and revealed individual- and genotype-specific patterns of ciliary dysfunction not captured by a single assay. Basolateral administration of *DNAI1* mRNA-loaded lipid nanoparticles resulted in partial, dose-dependent recovery of ciliary function in *DNAI1*-deficient HNECs.

**Conclusion:** This study establishes a standardized, individual-specific multi-assay nasal epithelial platform for functional phenotyping of motile cilia and preclinical evaluation of emerging therapies, with demonstrated utility in PCD.

## Introduction

Impaired motile cilia function is an underrecognized driver of chronic respiratory disease through loss of mucociliary clearance, a central airway defense mechanism against inhaled pathogens and environmental insults. This is exemplified by primary ciliary dyskinesia (PCD), a monogenic disorder caused by defects in motile cilia, resulting in impaired mucociliary clearance, chronic respiratory infections, and the development of bronchiectasis [1]. Although genetic and ultrastructural diagnostics have greatly improved disease classification, they do not fully capture the functional diversity of ciliary defects observed across individuals, as pathogenic variants in more than 50 genes can give rise to distinct structural and functional phenotypes [1]. As mutation-targeted and RNA-based therapeutic strategies begin to emerge [2], there is an increasing need for individual-specific epithelial systems that enable direct assessment of ciliary dysfunction and therapy responses.

In cystic fibrosis, nasal epithelial culture models have become increasingly important for linking genotype to epithelial function and for predicting response to cystic fibrosis transmembrane conductance regulator-modulating therapies [3–5]. A similar opportunity exists for motile ciliopathies, where direct functional assessment of motile cilia in nasal epithelia of patients could improve phenotypic stratification and support the development of emerging therapies [6]. However, current nasal epithelial cultures often exhibit variable multiciliogenesis, leading to heterogeneous cultures with limited reproducibility and thereby limiting robust quantitative assessment of ciliary phenotypes [7, 8]. Moreover, motile cilia dysfunction is inherently multidimensional and may manifest as defects in ciliogenesis, beat frequency, or coordinated motility. This complexity is reflected in the range of *in vitro* assays used to assess ciliary function, including ciliary beat frequency and mucociliary bead transport in 2D air–liquid interface cultures, and more recently in apical-out organoid based rotational readouts abnormalities [9–14]. Yet, a systematic framework that integrates these complementary assays in a standardized manner is currently lacking.

In this study, we established and validated a personalized multi-assay *in vitro* platform with primary human nasal epithelial cells (HNECs) for the assessment of motile ciliary dysfunction, using PCD as lead disease context. By optimizing the differentiation of HNECs toward a reproducibly multiciliated phenotype, we generated a standardized system compatible with complementary 2D and 3D assays of ciliary function. Using this framework, we captured genotype-specific ciliary dysfunction across a panel of PCD genotypes and assessed therapeutic rescue following delivery of *DNAI1* mRNA-loaded lipid nanoparticles (LNPs). Together, these findings position this platform as a scalable approach for individual-specific phenotyping and preclinical therapeutic testing for respiratory motile ciliopathies such as PCD.

## Material and methods

Please see the supplementary material for a detailed method section.

## Results

### Combined Notch and BMP inhibition, and apical medium promote ciliated cell differentiation

We first aimed to optimize the differentiation conditions for cryopreserved HNECs to generate highly ciliated cultures compatible with cilia function assays. This was accomplished by using our previously described differentiation medium containing small-molecules inhibiting the γ-secretase/Notch (DAPT) and BMP/SMAD (DMH-1) signaling pathways, which promotes ciliated cell differentiation in submerged cultures (**Figure 1A**) [15]. In HNECs from healthy controls, combined treatment with DAPT and DMH-1 showed a synergistic effect, significantly increasing immunofluorescence (IF) staining of β-tubulin IV compared to control conditions (**Figure 1B, C**). In contrast, the coverage of MUC5AC-positive secretory cells remained similar (**Figure 1B, D**). In addition, we investigated the effect of applying a thin apical liquid layer, which previously has been shown to increase ciliated cell differentiation [15, 16]. Combining low liquid-liquid interface conditions with DAPT and DMH-1 further increased ciliation while reducing differentiation towards secretory cells compared to control cultures. In contrast, addition of BMP4 increased secretory cell differentiation without affecting the ciliated area (**Figure 1B-D**). We further investigated the combined effects of DAPT, DMH-1, and low liquid-liquid interface conditions on mucociliary differentiation dynamics in HNECs from healthy individuals **(Supplementary Figure 1A**). Ciliated cell coverage was low at differentiation day 7 and increased and remained stable from day 14 onwards (**Supplementary Figure 1B, C**), whereas secretory cell area peaked at day 7 and decreased continuously thereafter (**Supplementary Figure 1B, D**). Analyses of mRNA expression levels of *MUC5AC* showed a similar pattern. In contrast, *FOXJ1* (ciliated cell marker) expression was detectable at low levels at day 7 and only increased significantly at day 42. Expression levels of *TP63* (basal cell marker) remained stable throughout differentiation (**Supplementary Figure 1E-G**). Altogether, our results demonstrate that dual Notch and BMP signaling inhibition by DAPT and DMH-1, combined with low liquid-liquid interface-conditions, lead to ciliated cell enrichment in nasal epithelial cultures.

**Figure 1.**
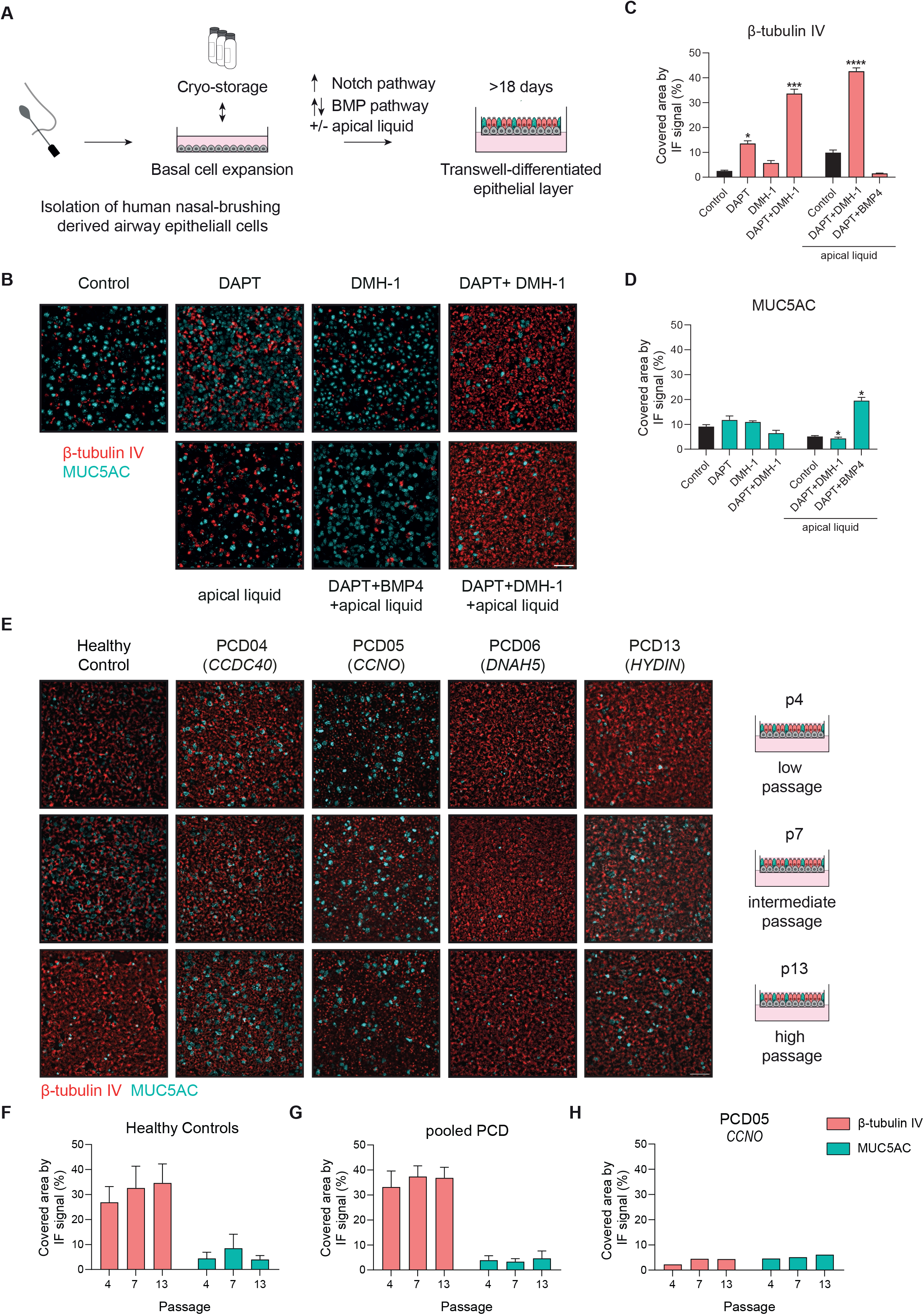
Differentiation of human nasal epithelial cells (HNECs) and induction of ciliated cell formation by small molecules and apical liquid, across low to high passages in healthy and primary ciliary dyskinesia (PCD) derived HNECs. **A** Graphical illustration of the workflow for expanding (cryo-stored) nasal brushing-derived airway basal cells and differentiating them in a transwell culture system to generate a differentiated epithelial monolayer. **B** Representative immunofluorescence (IF) images of an 18-day differentiated monolayer derived from HNECs of a healthy control using medium containing DAPT and/or DMH-1 or BMP4 without (air-liquid interface cultures) or with a small volume of apical medium (low liquid-liquid interface cultures). Cells were stained for the ciliated cell marker β-tubulin IV (red) and the secretory cell marker MUC5AC (cyan). **C** Quantification of β-tubulin IV and **D** MUC5AC IF staining. **E** Representative IF images of ciliated-cell marker β-tubulin IV (red) and secretory cell marker MUC5AC (cyan) of a healthy control and four PCD individuals with alterations in the *CCDC40, CCNO, DNAH5* and *HYDIN* genes at passage 4, 7, and 13 at 18 days post-differentiation. **F** Quantification of β-tubulin IV and MUC5AC signal measured in low liquid-liquid interface differentiated HNECs of healthy controls and **G, H** PCD-derived HNECs at passage 4, 7, 13. Healthy control data derived from three different individuals (left panel). PCD pooled data are composed of data from PCD04, PCD06 and PCD13 (middle panel). Data from PCD05 are plotted separately (right panel). Results are shown as mean ± SEM of HNECs from three individuals imaged at three randomly selected regions. Scale bars, 50 µm, * p < 0.05, *** p < 0.001, and **** p < 0.0001 as compared to control without apical liquid in panels **C, D** (Kruskal-Wallis test followed by a Dunn’s multiple comparison test), to passage 4 in panels **F, G** (one-way ANOVA with Dunnett’s multiple comparisons).

### Ciliated cell enrichment persists in HNECs after long-term expansion

Reduced differentiation of ciliated cells after long-term expansion is a common limitation of primary airway epithelial cells [17, 18]. Therefore, we characterized differentiation capacity of long-term expanded HNECs under ciliated cell enrichment conditions. This was evaluated by comparing differentiated nasal cell cultures at low (p4), intermediate (p7), and high passages (p13) at 18 days post-differentiation (**Figure 1E-H**). The capacity for differentiation into secretory and ciliated cells was maintained across passages, as indicated by stable MUC5AC and β-tubulin IV IF staining (**Figure 1E, F**). 2D differentiated HNECs derived from PCD individuals carrying *CCDC40, DNAH5*, or *HYDIN* pathogenic variants showed comparable secretory and ciliated cell areas to healthy controls (**Figure 1E, G**). In cultures carrying a *CCNO* pathogenic variant, ciliated area was lower than in healthy controls, consistent with previous research showing reduced numbers or absence of cilia [1, 19]. However, secretory cell area remained comparable to healthy controls (**Figure 1E, H**). Combined, these findings demonstrate that ciliated cell enrichment conditions support stable differentiation of both healthy and PCD-derived HNECs across low to high passages.

### Highly ciliated HNECs enable ciliary beat frequency and bead transport measurements

To further validate our ciliated cell enrichment conditions, we evaluated motile cilia function in HNECs from healthy controls at different time points during differentiation. First, we quantified the active ciliated area and ciliary beat frequency (CBF) using high-speed imaging (2D cilia activity, **Figure 2A-D**). The active ciliated area continuously increased over time (**Figure 2B, C**), whereas CBF was stable throughout differentiation (**Figure 2D)**. CBF was independent of active area (r^2^ = 0.1062; **Supplementary Figure 2**). Next, we assessed mucociliary transport using a bead transport (BT) assay in which 15μm polystyrene beads are displaced by ciliary activity across the epithelial surface (2D bead transport, **Figure 2E, F**). In contrast to single bead tracking approaches [10, 14], our method quantifies bead flow velocity and vorticity, a measurement of local rotational trajectory, providing a more comprehensive assessment of mucociliary transport. Both bead velocity and vorticity showed an increasing trend over time (**Figure 2G, H**), and were positively correlated (r^2^ = 0.5252, **Figure 2I**), indicating that faster BT is associated with more rotational trajectories. At day 25, bead movement often followed a distinct spiral trajectory **(Figure 2F, Supplementary Video 1**). Overall, these results demonstrate that our ciliated cell enrichment conditions promote a time-dependent increase in coordinated motile cilia activity in HNECs from healthy controls.

**Figure 2.**
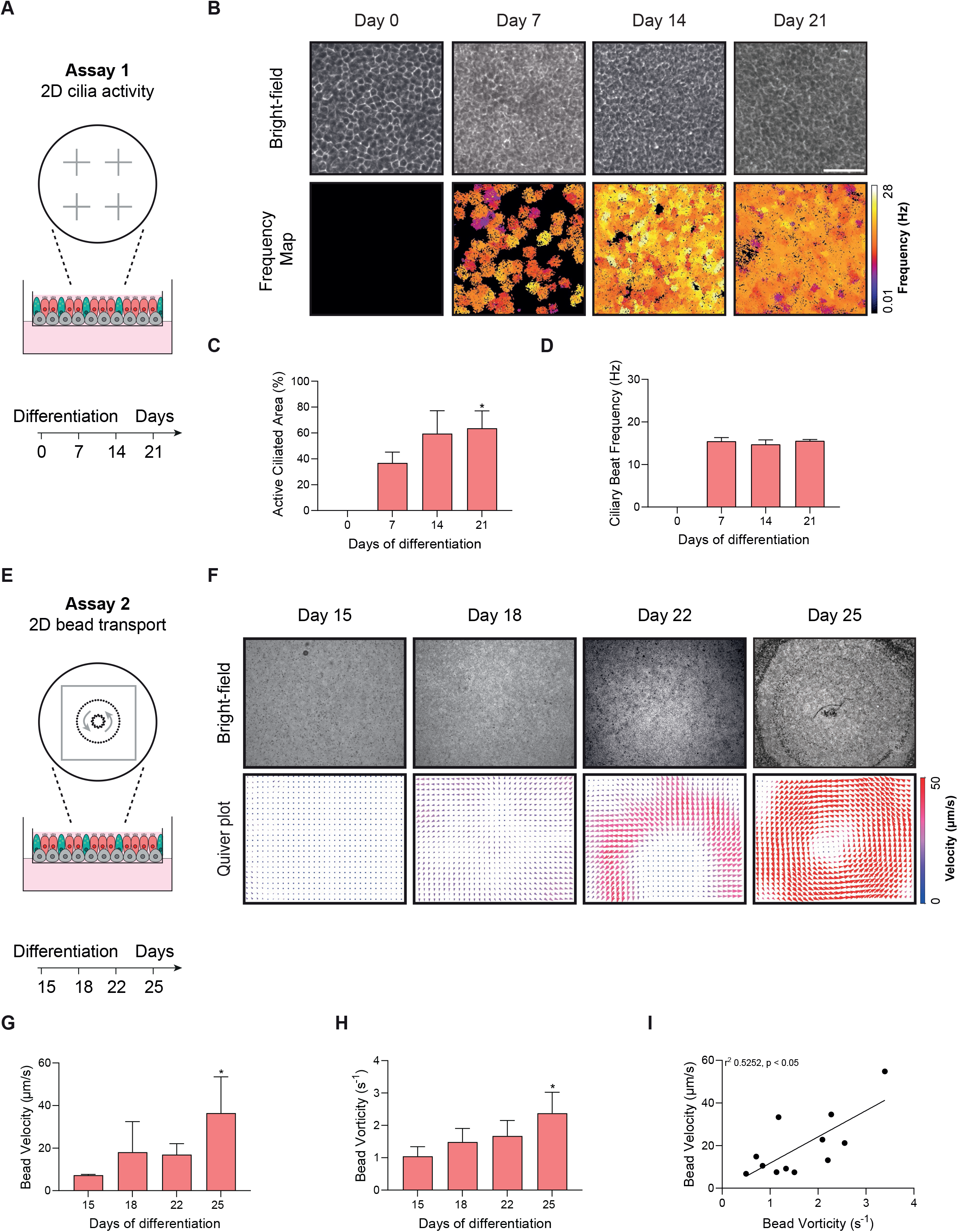
Cilia activity and bead transport in 2D-differentiated HNECs from healthy individuals over time. **A** Graphical illustration of the four positions and the different measurement timepoints used to validate 2D cilia activity (assay 1). **B** Representative bright-field video frames and corresponding frequency heatmaps of cilia enriched cultures from a healthy individual at days 0, 7, 14, and 21 post-differentiation, together with **C** quantification of active ciliated area (%) and **D** ciliary beat frequency (CBF, Hz) within this area. Color bar indicates CBF from 0.01 to 28 Hz. **E** Graphical illustration of the bead rotational movement and the different measurement timepoints to validate 2D bead transport (assay 2). **F** Representative bright-field video frames of 15 µm beads added to the apical surface of cultures from a healthy individual at days 15, 18, 22, and 25 post-differentiation. Corresponding quiver plots show bead velocity (µm/s) and direction (shown as arrows). Color bar indicates velocity from 0 to 50 µm/s. **G** Quantification of bead transport (BT) velocity (µm/s) and **H** vorticity (s^-1^) over time. **I** Correlation graph between bead vorticity and velocity (r^2^ = 0.5252). Results are shown as mean ± SEM of HNECs from three healthy individuals. Scale bars, 50 μm. * p < 0.05, ** p < 0.01, and *** p < 0.001, as compared to differentiation day 0 in panel **C, (**Friedman test with Dunn’s multiple comparisons), to differentiation day 7 in panel **D**, (Friedman test with Dunn’s multiple comparisons), to differentiation day 15 in panels **G, H** (Friedman test). In panel **I**, a Spearman correlation test was performed.

### Apical-out nasal organoid rotation as a readout of motile cilia function

In addition to 2D differentiated HNECs, we further characterize motile cilia function by measuring cilia-driven rotation of apical-out nasal organoid (AONOs; 3D cilia activity). In contrast to previously published methods [20, 21], these organoids were generated from fragmented 2D monolayers differentiated under ciliated cell enrichment conditions. Maintaining these epithelial fragments on an orbital shaker in the absence of extracellular matrix induces polarity inversion, resulting in structures with outward-facing cilia (**Figure 3A, Supplementary Videos 2, 3**). IF staining for β-tubulin IV verified apical-out orientation, with ciliary microtubules visible on the surface of ciliated cells in AONOs (**Figure 3B, Supplementary Figure 3A**). To quantify motile cilia function, a droplet of 4 μL Matrigel with at least 100 AONOs was dispensed in each well of a 96 well-plate (**Supplementary Figure 3B-C**). First, all organoids within each droplet were segmented using OrgaSegment [22] and subsequently their angular speed was analyzed using an organoid rotation analysis model [20] (**Figure 3C**). Using this approach, we observed that the percentage of rotating organoids was the highest when embedded in 50% Matrigel after 96 hours on an orbital shaker (**Supplementary Figure 3D, E**). Therefore, this condition was used as the standard approach. To validate the rotation assay, we assessed the effect of the motile cilia inhibitor paclitaxel [23, 24], which induced a dose- and time-dependent reduction in the percentage of rotating AONOs (**Figure 3D, E**). In addition, paclitaxel significantly decreased cell viability, which may have contributed to this observed decrease in organoid rotation (**Supplementary Figure 3F**).

**Figure 3.**
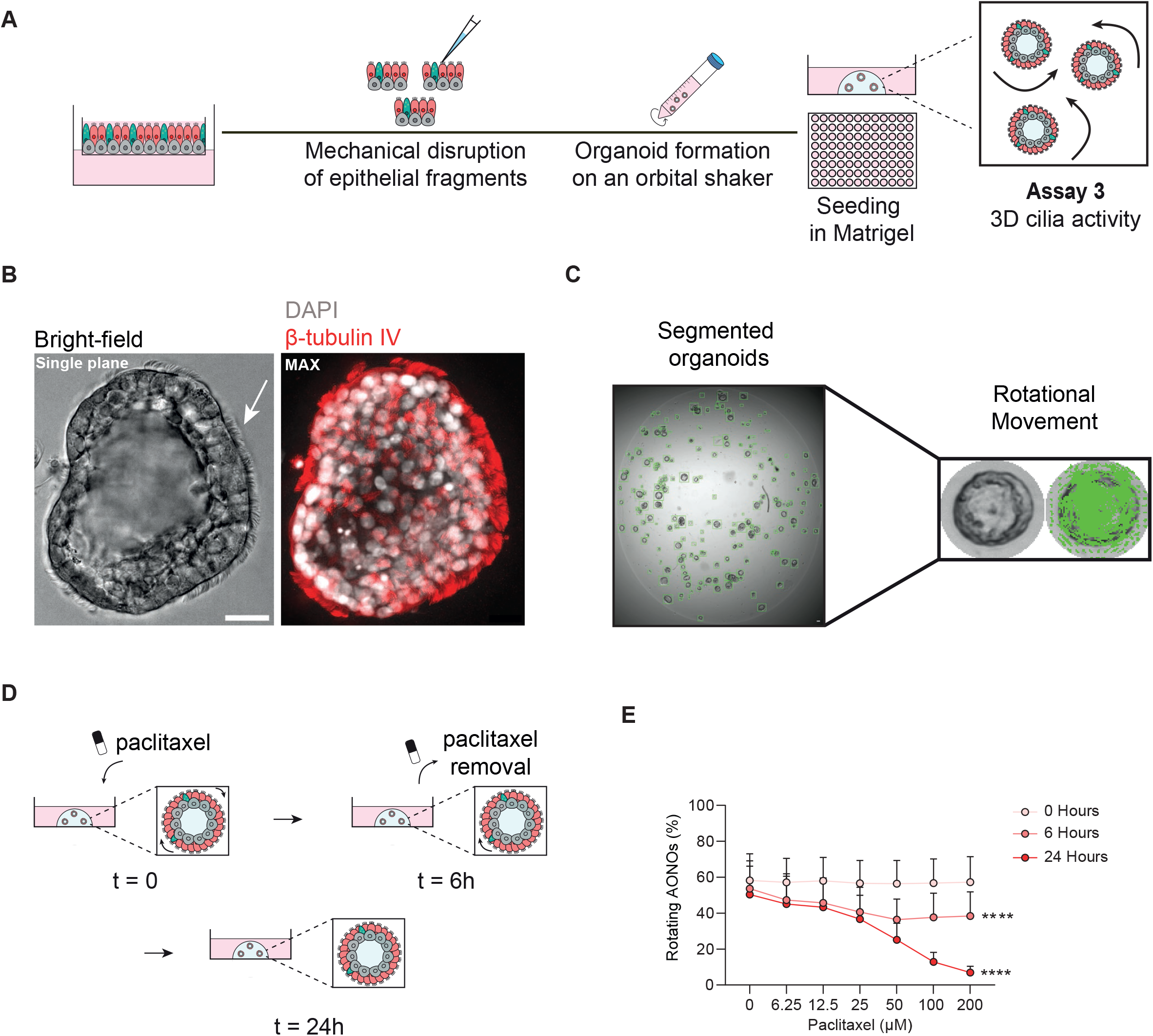
3D rotation of apical out nasal organoids (AONOs) derived from differentiated HNECs of healthy individuals. **A** Graphical illustration showing the procedure from disrupting the differentiated monolayer to seeding of the AONOs in a 96-well plate. **B** Representative bright-field image of an AONO derived from a healthy individual, showing cilia facing outwards (indicated with white arrow) and stained for the nuclear marker DAPI (gray) and the ciliated cell marker β-tubulin IV (red). Scale bar, 25 μm. **C** Representative image of segmented AONOs including indication of rotational movement of an individual organoid. Scale bar, 50 μm. **D** Schematic overview of the experimental workflow of the addition of paclitaxel on AONOs and **E** quantification of the percentage of rotating AONOs generated from healthy individuals treated with various concentrations of paclitaxel at t = 0h (before treatment), t = 6h (immediately after treatment removal), and t = 24h (18h after treatment removal). Green lines indicate rotational movement. Results are shown as mean ± SEM of HNECs from one healthy individual. **** p < 0.0001 in panel **B** (linear mixed model followed by a type III ANOVA).

Overall, the AONO rotation assay enables simultaneous quantification of both the number and angular speed of hundreds of AONOs per Matrigel droplet in a 96 well plate format, thereby providing a scalable platform for functional assessment and therapeutic screening.

### Three functional ciliary assays capture different aspects of ciliary dysfunction

Next, we determined motile cilia dysfunction in differentiated HNECs from individuals with PCD using the established functional ciliary assays. Both 2D and 3D assays revealed differences in ciliary function between PCD and healthy control samples (**Figure 4A, Supplementary Figure 4A-F**), which were further highlighted using z-score normalization relative to healthy control (**Figure 4B, Supplementary Figure 5A**). To examine how the different functional readouts relate to each other, we used assay-level correlation matrices across all individual HNEC samples and within the PCD group separately. When including all samples, most assays showed strong correlations (**Figure 5B**). In contrast, within the PCD group, correlations between different functional layers were generally weak. However, correlations were observed within assays measuring the same functional layer, including frequency and active beating area, bead velocity and vorticity, and AONO angular speed and rotation percentage (**Figure 5C**). Based on this, we combined related outcome parameters into composite block scores representing 2D cilia activity (assay 1), 2D bead transport (assay 2), and 3D cilia activity (assay 3). Boxplots of these block scores showed clear separation between healthy control and PCD groups (**Supplementary Figure 5B-D**). This underlined that correlations seen in the full cohort are partly driven by differences between PCD and healthy control (**Figure D-E**). Within the PCD group, block scores were weakly correlated, indicating that ciliary activity, bead transport, and 3D function capture partly, independent aspects of ciliary dysfunction (**Figure 5E**). Consistent with this, scatter plots of PCD samples revealed donor-to-donor heterogeneity. Some genotypes showed a more restricted distribution in ciliary activity versus BT or versus rotation, suggesting genotype-associated patterns (**Figure 5F, G**). For example, variants such as *DNAH11* and *HYDIN* showed preserved motility despite structural abnormalities [9, 19]. However, this pattern was less apparent when comparing transport and rotation, where individuals carrying the same variant showed divergent behavior (**Supplementary Figure 5E**), as illustrated by distinct functional phenotypes observed in different *CCDC39* variants. Together, these multi-assay ciliary function analyses revealed PCD individual-specific patterns of ciliary dysfunction that cannot be captured by a single functional readout.

**Figure 4.**
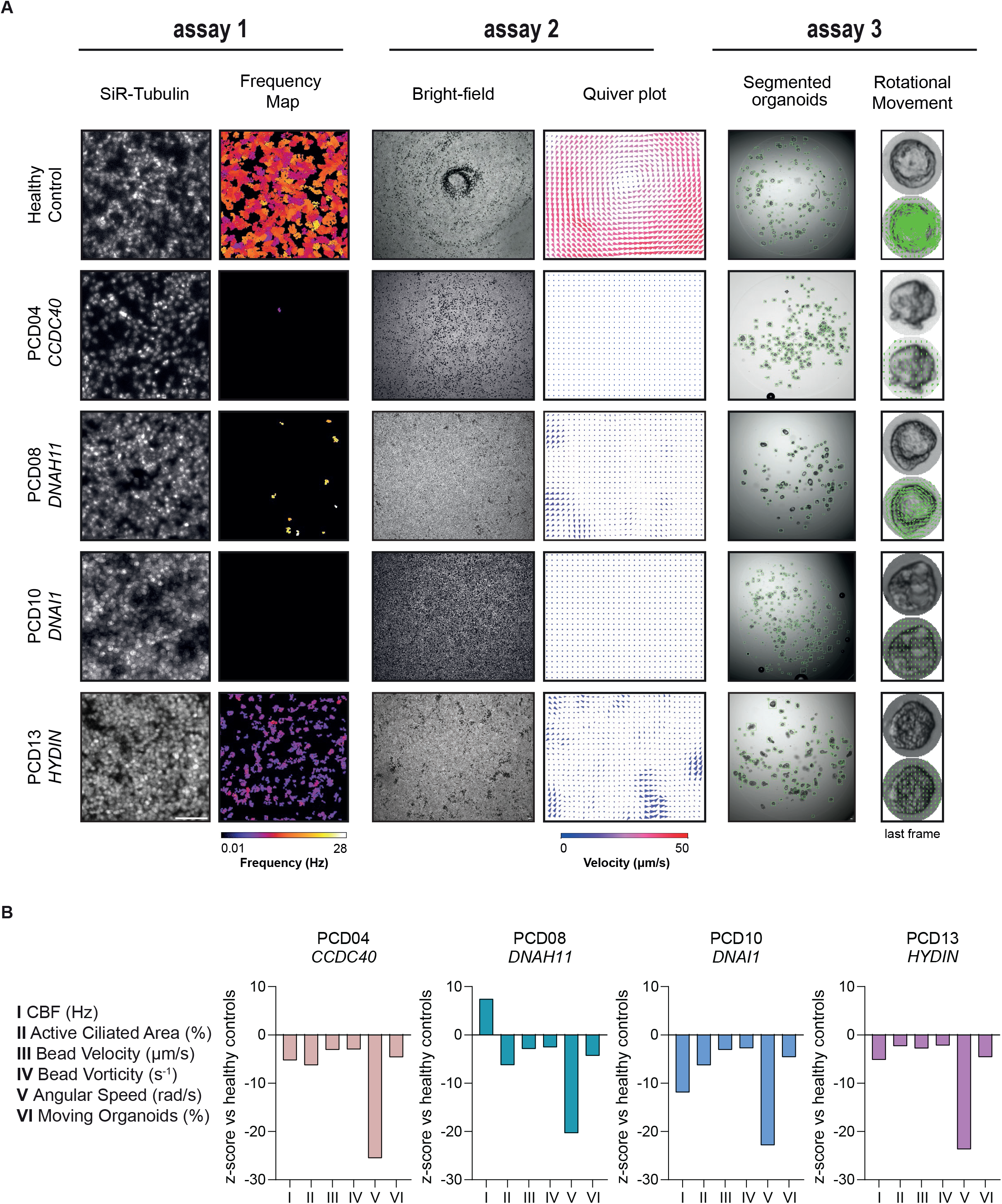
An *in vitro* multi-assay platform distinguishes between healthy and PCD defects. **A** The *in vitro* multi-assay platform applied on 2D differentiated HNECs of a healthy individual and four PCD individuals with a genetic variant in the *CCDC40* (PCD04), *DNAH11* (PCD08), *DNAI1* (PCD10), or *HYDIN* (PCD10) gene. Representative images of HNECs stained with a live cilia staining marker SiR-Tubulin and the corresponding frequency heatmaps. Color bar indicates CBF from 0.01 to 28 Hz (assay 1). Bright-field video frames depicting 15 µm beads added apically and the corresponding quiver plots. Color bar indicates velocity from 0 to 50 µm/s (assay 2). Bright-field video frames of segmented AONOs and motion-tracking visualization of individual organoids. Green lines indicate rotational movement (assay 3). **B** Z-score of each ciliary functional measurement for HNECs with a genetic alteration in the *CCDC40* (PCD04), *DNAH11* (PCD08), *DNAI1* (PCD10), or *HYDIN* (PCD10) genes normalized to healthy controls. Mean and SD were calculated from the healthy controls, and z-score for each measurement was calculated relative to the healthy control mean. Scale bars, 50 µm.

**Figure 5.**
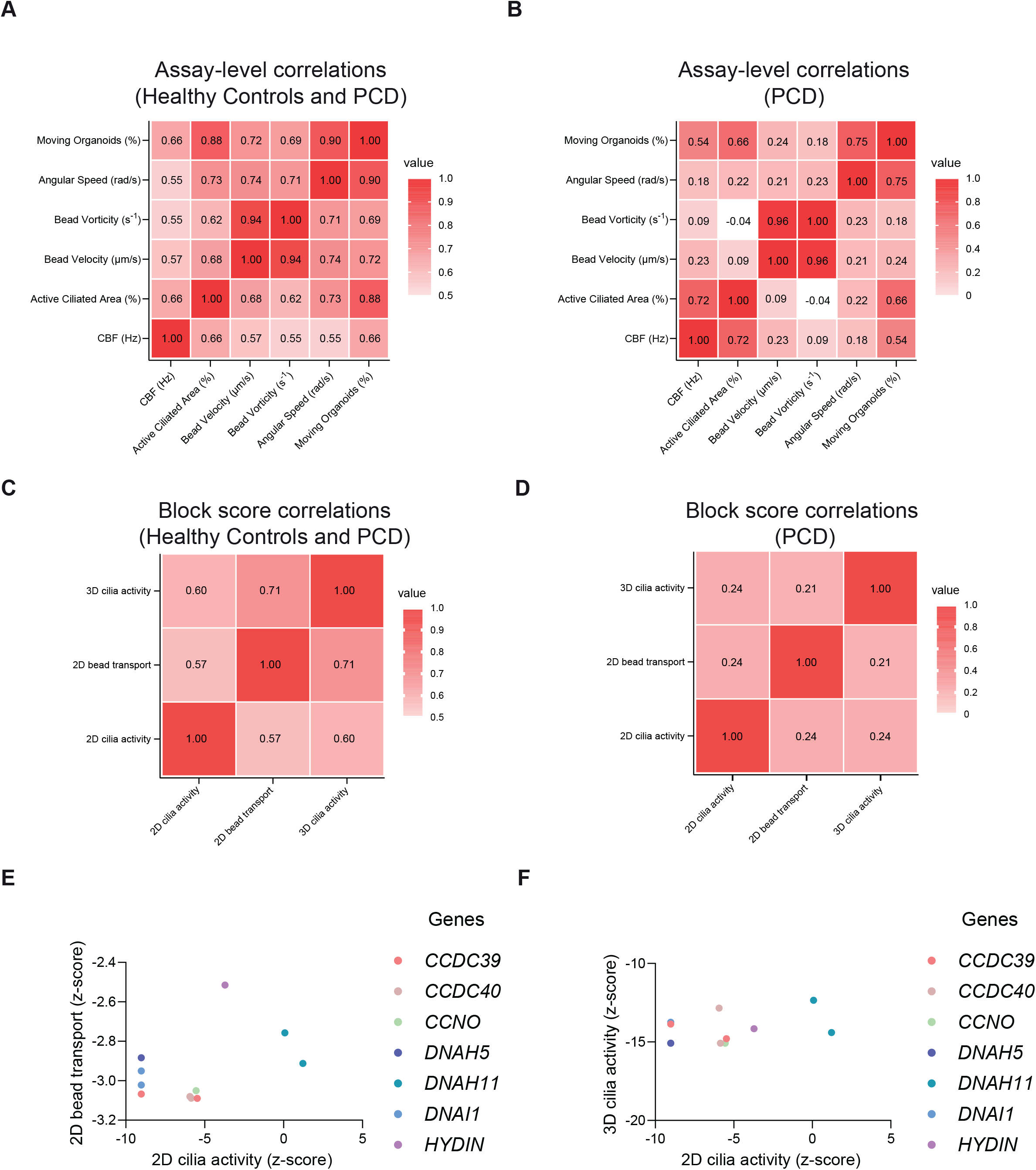
Integrated analysis of ciliary activity assays distinguishes healthy and PCD samples. **A** Correlation matrice of z-score normalized measurements across the indicated cilia functional measurements for five healthy and 11 PCD individuals and **B** for solely the 11 PCD individuals. **C** Correlation matrice of assay block scores including 2D activity, 2D transport and 3D rotation for five healthy and 11 PCD individuals and **D** for solely the 11 PCD individuals. **E** Correlation graph between 2D cilia activity (assay 1) and 2D BT (assay 2) and **F** between 2D cilia activity (assay 1) and 3D cilia activity (assay 3) shown as z-score values for the 7 different PCD genes. For panels **A-D**, pairwise correlations between variables were computed using Spearman correlation coefficients. Color bar indicates the correlation coefficient values. For panels **C, D**, block scores were calculated as the mean of the z-scores of the two parameters corresponding to each assay: CBF with active ciliated area corresponds to the 2D activity, bead velocity with vorticity to 2D bead transport and angular speed with moving organoids to 3D rotation.

### Rescue of DNAI1 deficiency using mRNA-loaded lipid nanoparticles

To validate our 2D differentiated HNEC model in a therapeutic context, we examined ciliary recovery in cells from PCD subjects with *DNAI1* genetic alterations after treatment with *DNAI1* mRNA-loaded LNPs. We first examined the uptake of LNPs in differentiated HNECs using a *tdTomat*o mRNA reporter. Following a single basolateral treatment, uptake was higher in HNECs cultured on transwell inserts with a 1.0 μm pore size compared to 0.4 μm (**Supplementary Figure 6A, B**). Moreover, active ciliated area and frequency were comparable between HNECs differentiated on both filter types (**Supplementary Figure 6C, D**). Based on these findings, all subsequent experiments evaluating ciliary recovery were performed using 1.0 μm pore size inserts. Ciliary recovery was assessed in HNECs from two independent PCD individuals carrying *DNAI1* genetic variants using previously published *DNAI1* mRNA-LNPs [25, 26]. To evaluate sustained therapeutic rescue, basolateral LNP treatments were administered three times a week starting at day 14 post differentiation (**Figure 6A**). Restoration of DNAI1 protein expression was confirmed by western blot and by IF staining, where DNAI1 localization was observed in close proximity to the β-tubulin IV-positive ciliated regions. Overall, protein expression was increased in treated conditions compared to untreated *DNAI1*-deficient cultures, although levels did not reach those observed in healthy controls (**Figure 6B, C**). Cilia recovery was also confirmed on a functional level with mRNA-LNP treatments resulting in a dose-dependent increase in the percentage of ciliary active area, reaching approximately one-third of that observed in healthy controls (**Figure 7A, B**). Across all treatment conditions, the CBF remained comparable (**Figure 7C**). Consistent with these findings, the BT assay revealed a dose-dependent increase in cilia-driven particle movement, also reaching approximately one-third of the velocity measured in healthy controls (**Figure 7D, E**). Vorticity showed a similar dose-dependent increase and reached approximately two-thirds of healthy control levels (**Figure 7F**). Bead velocity strongly correlated with vorticity in DNAI1-PCD derived cells treated with LNP-mRNA (r^2^ = 0.8924; **Supplementary Figure 7A**). In addition, active ciliated area showed moderate-to-strong correlations with bead velocity (r^2^ = 0.6921) and vorticity (r^2^ = 0.7574), indicating that increased ciliated cell coverage is associated with improved BT parameters (**Supplementary Figure 7B, C**). Together, these findings demonstrate that mRNA-based LNP treatment partially restores ciliary function in *DNAI1*-deficient HNECs at both the protein and the functional level.

**Figure 6.**
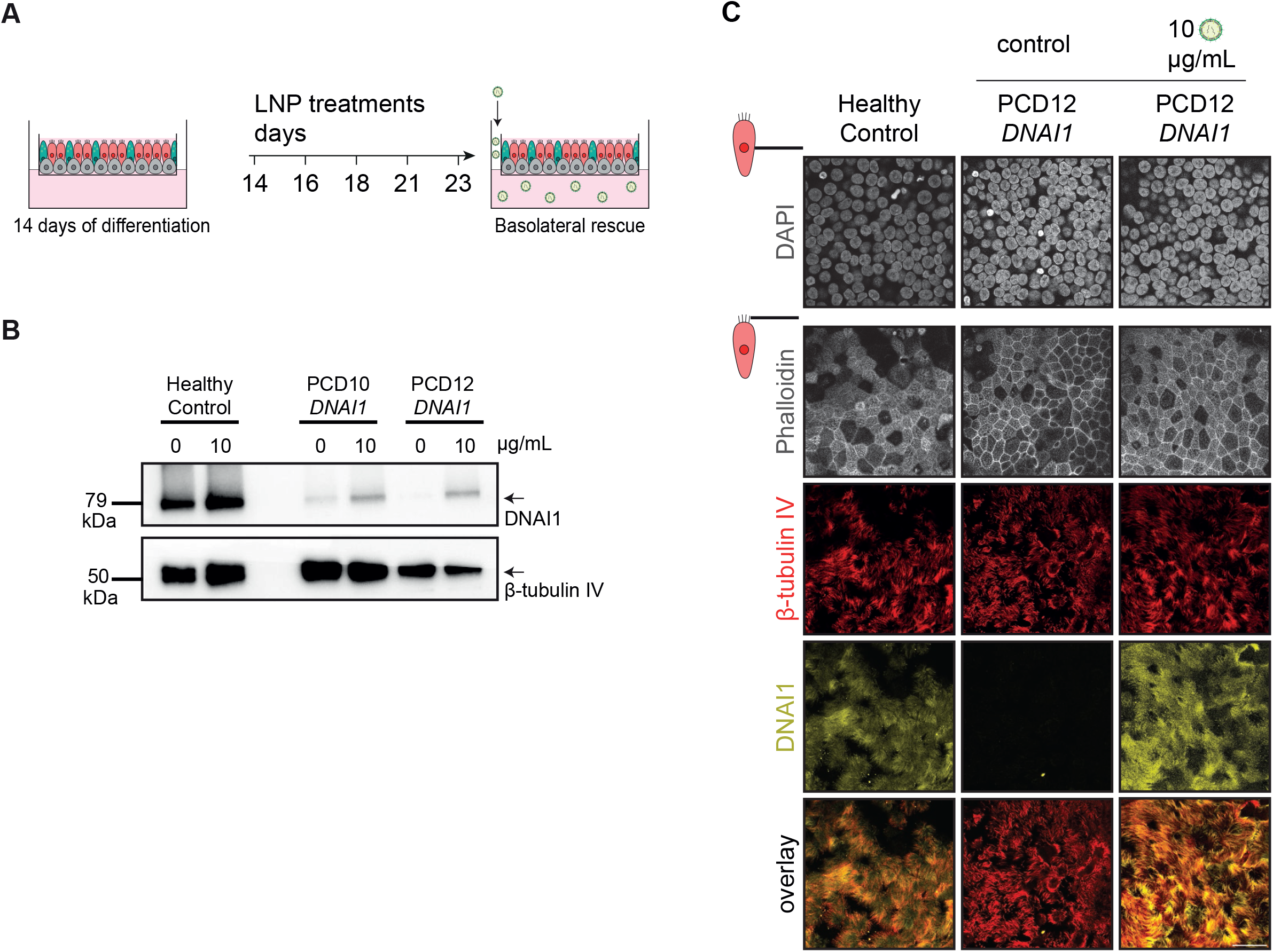
Therapeutic treatment of *DNAI1* mRNA delivery via lipid nanoparticles (LNPs) partially rescues DNAI1 protein expression in PCD-derived HNECs. **A** Graphical illustration of mRNA delivery in 2D differentiated HNECs treated with five basolateral LNP treatments from day 14 to 23 post differentiation. **B** Cropped western blots showing DNAI1 protein expression with β-tubulin IV as an internal control in a healthy individual and two PCD individuals with *DNAI1* alterations (PCD10 and PCD12) treated with or without 10 µg/mL *DNAI1* mRNA LNPs. **C** Representative IF stainings of 2D cultures from a healthy control and a PCD sample (PCD12) treated with or without 10 µg/mL *DNAI1* mRNA LNPs. Cells were stained for the nuclear marker DAPI (gray), filamentous actin marker phalloidin (gray), ciliated cell marker β-tubulin IV (red), and DNAI1 (yellow), imaged at two layers (top and bottom panel). Scale bar, 25 µm

**Figure 7.**
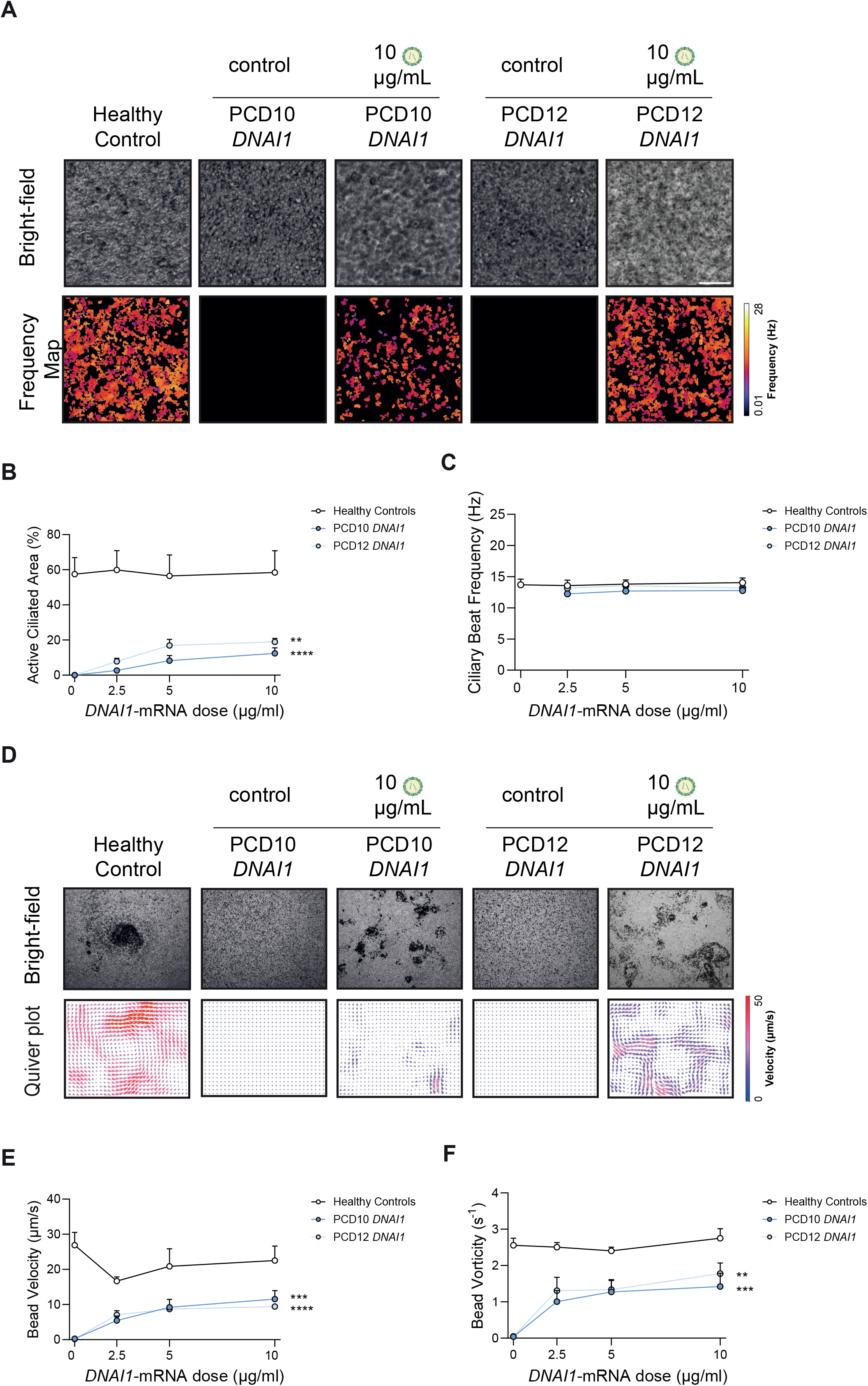
2D mucociliary clearance rescue of DNAI1 alterations using *DNAI1* mRNA delivery via lipid nanoparticles. **A** Representative bright-field video frames with corresponding frequency heatmaps of 2D differentiated HNECs from a healthy untreated individual and two PCD individuals with *DNAI1* genetic alterations treated with or without 10 µg/mL *DNAI1* mRNA LNPs. Color bar indicates CBF from 0.01 to 28 Hz. **B** Quantification of the percentage of active ciliated area and **C** CBF within these active areas of HNECs from healthy and PCD individuals treated with different *DNAI1* mRNA LNP concentrations. **D** Representative bright-field video frames of 15 µm beads added to the apical surface of 2D cultures from a healthy individual and two PCD individuals carrying *DNAI1* genetic alterations at day 25 post differentiation, treated with or without 10 µg/mL *DNAI1* mRNA LNPs. Corresponding quiver plots show bead velocity (µm/s) and direction (shown as arrows). Color bar indicates velocity from 0 to 50 µm/s. **E** Quantification of bead transport velocity and **F** vorticity of 2D cultures from healthy and PCD individuals treated with different concentrations of *DNAI1* mRNA LNPs. Scale bars, 50 μm. Results are shown as mean ± SEM of HNECs from two pooled healthy individuals and two PCD individuals with a *DNAI1* alteration. In panels **B, C, E** and **F**, data from 2D-differentiated HNECs of healthy controls were fitted with a linear regression and of DNAI1 individuals with a non-linear one-phase association curve (nested-model ANOVA).

## Discussion

In the present study, we developed and validated a personalized multi-assay platform to assess motile ciliary function in HNECs, using PCD as a representative disease context (**Figure 8**). This approach enabled genotype-specific functional profiling of ciliary dysfunction across 11 distinct PCD genotypes and supported proof-of-concept therapeutic evaluation using *DNAI1*-loaded LNPs.

**Figure 8.**
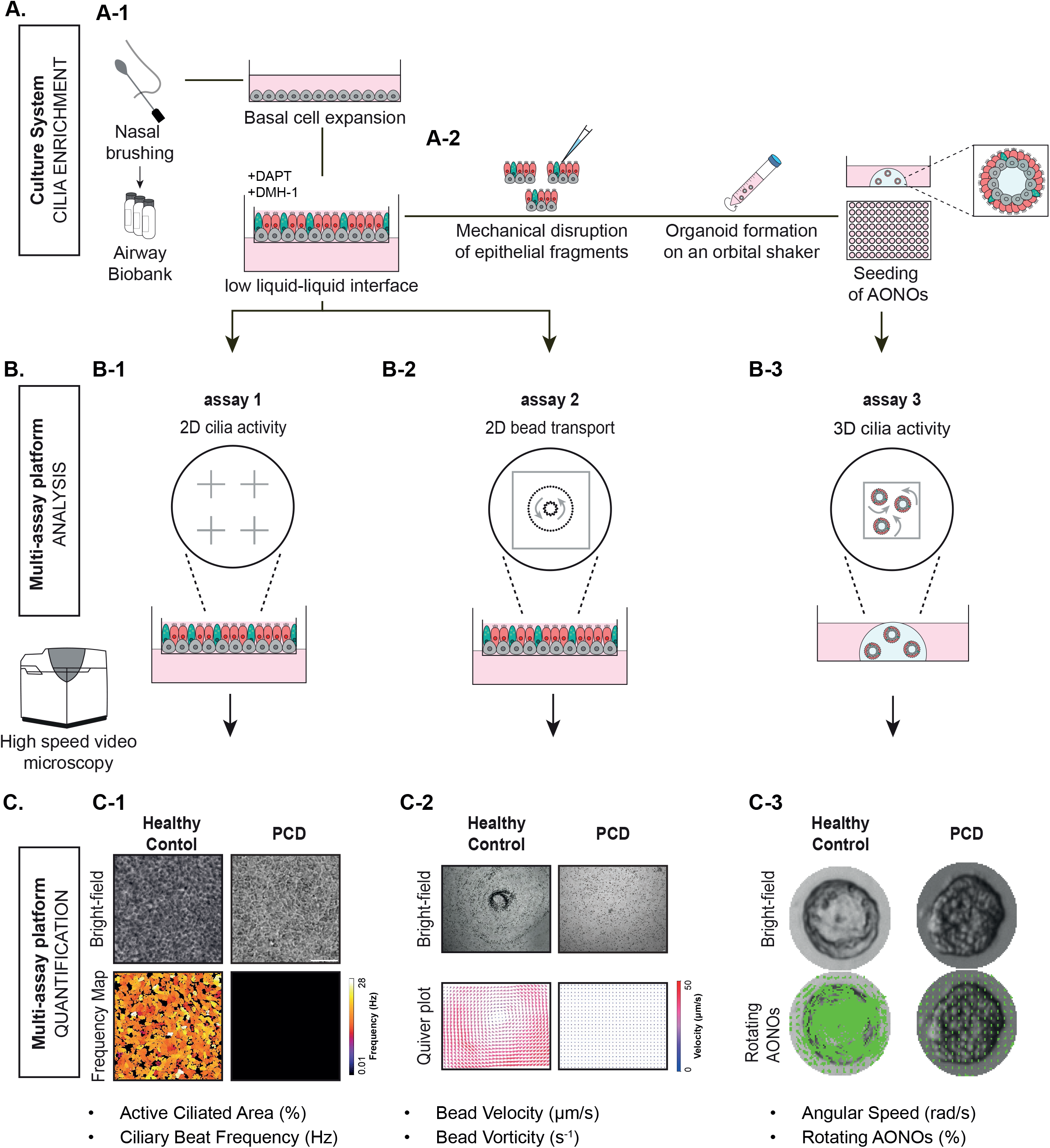
A personalized multi-assay *in vitro* platform to assess ciliary (dys)function. **A-1** Graphical illustration of the workflow for expanding (cryo-stored) nasal brushing-derived HNECs and 2D differentiation of ciliated cell-enriched cultures. **B** Workflow of the multi-assay platform integrating three complementary ciliary function assays: 2D cilia activity (assay 1), 2D BT (assay 2), and 3D cilia activity (assay 3). **B-1** CBF (assay 1) was measured using high-speed video microscopy at four positions per well, imaged at 5x magnification (bright-field). **C-1** Quantification was performed by generating frequency maps showing the percentage of active ciliated area and the beat frequency (Hz) within this active area. **B-2** For the BT assay (assay 2), 15 µm polystyrene beads were added to the apical surface of epithelial cells and bead movement was recorded at 2.5x magnification for 30 seconds per well (bright-field). Videos were captured at or near the center of each well. **C-2** Quantification was performed using particle image velocimetry, showing bead velocity (µm/s; blue to red for low to high velocity), direction (shown as arrows), and local rotation (vorticity, s^-1^). **A-2** For AONO generation (assay 3), 2D differentiated cultures were mechanically fragmented, continuously rotated for 1-4 days, and embedded in Matrigel droplets in 96-well plates to generate organoids with cilia facing outwards. **B-3** AONO rotation was recorded using 2.5x magnification for 30 seconds (bright-field). **C-3** Segmented organoid quantification of percentage rotating organoids and their angular speed (rad/s). These assays were performed on epithelial cells from the same wells, enabling direct correlation of functional parameters. Assays were performed on epithelial cells derived from both healthy and PCD individuals. Scale bars, 50 µm.

We demonstrated a synergistic increase in ciliogenesis when both Notch and BMP signaling pathways were inhibited, consistent with earlier observations [15]. Additionally, the presence of a thin apical liquid layer further promoted ciliated cell differentiation, although the underlying mechanism remains unclear [16]. Our culture conditions demonstrated an accelerated onset of cilia formation compared with previous reports, with a substantially higher active ciliated area at early time points [27, 28]. Moreover, ciliated cell differentiation was preserved in serial passaged HNECs, whereas previous studies report successful differentiation with limited passaging [29–32]. These findings indicate that HNECs can be expanded and repeatedly used while retaining their capacity for multiciliated differentiation, supporting their suitability for scalable and longitudinal functional studies.

By combining three complementary assays, we provide a comprehensive assessment of ciliary function in differentiated HNECs. Each assay offers unique insights, capturing beat frequency, coordinated mucociliary transport, or 3D dynamics, that are not apparent from a single readout. Indeed, some previous reports show CBF being measured in combination with bead transport. However, previous bead-tracking studies were limited by small imaging fields, high magnification requirements, reliance on fluorescent microbeads, and/or low throughput, often restricting analysis to localized regions rather than the epithelial surface as a whole [10, 13, 14, 33, 34]. Therefore, we developed a bright-field-based, low-magnification imaging setup combined with particle image velocimetry (PIV)-based analysis to enable scalable quantifications of bead movement across large regions of epithelial surface. Furthermore, in contrast to existing protocols [20, 35], our approach derives AONOs directly from 2D differentiated HNECs, enabling direct comparison between AONO rotation, CBF, and bead transport measurements. This multi-assay approach clearly distinguished healthy from PCD-derived HNECs. However, weak correlations between assay block scores within the PCD group indicate that these readouts capture independent aspects of dysfunction. Consistent with this, donor-to-donor heterogeneity was observed, suggesting genotype-associated patterns. Notably, bead transport parameters broadly aligned with known genotype-phenotype relationships in PCD, with variants such as *CCDC39, CCDC40*, and *CCNO* showing minimal transport, whereas *DNAH11* and *HYDIN* variants retained measurable activity, consistent with their comparatively milder clinical phenotypes [1]. Together, these findings demonstrate that PCD is characterized by multidimensional, individual-specific ciliary dysfunction that cannot be captured by a single functional assay. More broadly, this underscores the importance of integrated functional readouts to capture epithelial dynamics in complex airway diseases.

To explore the feasibility of using our culture model for testing RNA-based therapeutics, we investigated the administration of *DNAI1* mRNA-loaded LNPs from ReCode therapeutics [36] to HNECs harboring a *DNAI1* alteration. This resulted in partial recovery of ciliary function, which was captured by 2D ciliary function assays. Importantly, restoration of ciliary activity was accompanied by improved mucociliary transport, indicating that recovery of beating translates into functional clearance capacity in patient-derived epithelial cultures. The observed recovery, approximately one-to two-thirds of control levels, may already be sufficient to support functional mucociliary clearance, as previous studies have shown that 30–62% of functional multiciliated cells can be sufficient to produce normal or only mildly impaired mucociliary clearance [37]. Nevertheless, further optimization of dosing and delivery will likely be required to enhance therapeutic efficacy.

This study has several limitations. First, although the current cohort captures multiple mechanistic classes of PCD-associated dysfunction, it remains limited in size relative to the marked genetic heterogeneity of PCD [1]. Expanding the number and diversity of PCD subject-derived HNECs will be important to better resolve both intergenic and intragenic variability. Second, we did not directly correlate *in vitro* phenotypes with clinical *in vivo* outcomes. Although such comparisons remain uncommon in the PCD field, establishing links between epithelial functional readouts and individual-level measures of mucociliary clearance or disease severity will be important for further defining the translational relevance of these assays [37]. Third, our proof-of-concept mRNA-LNP experiments were limited to the basolateral compartment, whereas future studies should incorporate apical aerosol-based delivery and extend rescue assessment to 3D functional readouts. Finally, while this platform was developed and validated in the context of motile ciliopathies, its applicability to other respiratory diseases characterized by epithelial dysfunction, such as cystic fibrosis or chronic airway inflammation, remains to be established and needs further investigation.

Together, these findings demonstrate that standardized multiciliated nasal epithelial cultures enable individual-specific functional profiling of motile cilia in human airway epithelium. By integrating complementary 2D and 3D assays, this approach captures the multidimensional nature of ciliary dysfunction and provides a practical framework for evaluating emerging therapies in motile ciliopathies and potentially other airway diseases.

## Abbreviation list

AONO: Apical-Out Nasal Organoid
BT: Bead Transport
CBF: Ciliary Beat Frequency
HNEC: Human Nasal Epithelial Cell
IF: Immunofluorescence
LNP: Lipid Nanoparticle
PCD: Primary Ciliary Dyskinesia
PIV: Particle Image Velocimetry

## Acknowledgements

We thank the participants for donating nasal epithelial samples for this study. We acknowledge the Utrecht Platform for Organoid Technology (UPORT) for coordinating sample collection and biobanking.

