## Supplementary files for "Personalized multi-assay profiling of respiratory motile ciliopathies and mRNA therapy"

**Materials and methods**

**Human materials and sample collection**

Nasal brushings were collected from healthy volunteers without respiratory tract symptoms (n = 8 independent donors) and from individuals with PCD (n = 13 independent donors) as previously described [1]. All participants provided broad informed consent for the use of their samples in research. The study was reviewed and approved by the Biobank Research Ethics Committee (Toetsingscommissie Biobank Utrecht, the Netherlands) of the University Medical Center Utrecht (protocol ID: 16/586 and 25U-0676). The collection, storage, and distribution of nasal epithelial materials were coordinated by the Utrecht Platform for Organoid Technology, following standardized procedures and ethical oversight. PCD individual characteristics provided by Amsterdam Medical Centre and University Medical Center Utrecht includes affected gene, cDNA and protein changes, nasal nitric oxide levels, high speed video microscopy results of ciliary movement *ex vivo* before cell culture, and transmission electron microscopy (**Supplementary Table 1**).

**Isolation, expansion, and 2D differentiation of basal human nasal epithelial cells**

HNECs were isolated as previously described with minor modifications [1, 2]. Briefly, dissociated single cells were seeded onto collagen IV-coated plates (50 µg/mL; Sigma-Aldrich, St.Louis, USA) and cultured in isolation medium (**Supplementary Table 2**) containing amphotericin B, gentamicin, and vancomycin to prevent microbial contamination. After one week, medium was replaced with expansion medium supplemented with DAPT and rapamycin (**Supplementary Table 2**). HNECs were cultured to 80-90% confluency, with medium refreshed three times per week. Cells were maintained at 37°C in a 5% CO_2_ incubator and passaged using TrypLE^TM^ Express (Gibco, Grand Island, NY, USA). Differentiation of HNECs was performed as previously described with minor modifications [1]. Briefly, 0.2x10^6^ HNECs were seeded onto 24-well transwell inserts (0.4 μm pore size polyester membrane and 6.5 mm Inserts, Corning HTS, Corning, NY, USA) pre-coated with 30 µg/mL PureCol (Advanced BioMatrix, Carlsbad, CA, USA) and cultured in expansion medium (200 μL apical and 800 μL basolateral) until confluency. The medium was then replaced with differentiation medium supplemented with 500 nM A83-01 (**Supplementary Table 3**). Following formation of a tight epithelial monolayer, apical and basolateral medium volumes were reduced to 45 μL and 600 μL, respectively, establishing a low liquid-liquid interface culture. After 3-5 days, cultures were switched to ciliated cell enrichment medium, consisting of differentiation medium supplemented with 5 µM DAPT and 0.5 µM DMH-1. Medium was refreshed three times per week on the apical surface and twice per week on the basolateral side, and cultures were washed once per week using 100 µL PBS for 5 minutes at 37°C.

**Ciliary beat frequency assay to measure 2D ciliary activity (assay 1)**

CBF was measured in 2D differentiated HNECs using a ZEISS Celldiscoverer 7 microscope (ZEISS, Oberkochen, Germany) equipped with environmental control (37°C and 5% CO_2_). Four positions per transwell were imaged at 5x magnification, with a 288x288 pixel region of interest and an acquisition rate of 216 frames/s. CBF was quantified using the "Temporal ICS" command of the Correlescence v.0.0.5 plugin in ImageJ, based on a previously described analysis script and implemented here in a semi-automated analysis platform [3]. Briefly, the static component was removed by subtracting the average intensity image from the recorded time-lapse. For each pixel, the normalized autocorrelation function was calculated over time, and the first maximum (tolerance > 0.2) was used to determine the oscillation frequency. This generated a frequency map with the same X, Y dimensions as the original image, assigning a frequency value to each pixel. Areas with pixel sizes below 25 (representing subcellular background noise) were excluded from analysis. In parallel with CBF measurements, ciliary presence was confirmed by live SiR-Tubulin staining (Spirochrome, Stein am Rhein, Switzerland), a microtubule-labeling compound. For each CBF position, bright-field and SiR-Tubulin images were acquired immediately after CBF recordings. SiR-Tubulin was diluted 1:5000 in differentiation medium and applied to the apical (100 µL) and basolateral (600 µL) compartments for 4 h. After incubation, the staining medium was replaced with ciliated cell enrichment medium.

**Bead transport assay to measure 2D bead transport (assay 2)**

To quantify mucociliary transport in 2D differentiated HNECs, 15 μm polystyrene beads (Sigma-Aldrich; 10 % solids) were diluted 1:400 in differentiation medium, and 100 µL was added to the apical surface of the epithelial monolayer. Bead movement was recorded at 2.5x magnification using a ZEISS Celldiscoverer 7 microscope with a 2048x1640 pixel region of interest for 30 s/well, acquiring 1 frame/s. Videos were acquired from the center of each well and analyzed using PIV [4]. Prior to analysis, videos were preprocessed using the BackgroundSubtractorMOG2 algorithm implemented in OpenCV [5, 6], followed by erosion (2x2) and dilation (15x15) operations. Two frame sizes (256x256 and 128x128 pixels) were applied simultaneously during the PIV analysis to capture both small and large bead displacements (<https://github.com/Living-Technologies/cilliary_function_analysis>). Bead velocity (µm/s) was validated using single-bead tracking as a human-validated ground truth with Trackmate in ImageJ (data not shown). In addition to average bead velocity, flow coordination was quantified using the average absolute vorticity. Vorticity (ω), describing local rotational motion at a single point in the flow, was calculated as follows:

$\omega=\frac{\Delta V}{\Delta x}-\frac{\Delta U}{\Delta y}$

Where ΔV and ΔU represent changes in vertical and horizontal velocity, respectively, and Δy and Δx represent changes in vertical and horizontal distances, respectively.

Each frame was divided into equal square interrogation regions. Each square is represented by an arrow in the quiver plot and indexed by its spatial position i and j in the horizontal and vertical direction, respectively. The central differences between horizontally and vertically aligning blocks (divided over 2Δx and 2Δy, respectively) were calculated as follows:

$$\omega_{i,j}=\frac{V_{i+1,j}-V_{i-1,j}}{2\Delta x}-\frac{U_{i,j+1}-U_{i,j-1}}{2\Delta y}$$

Positive vorticity corresponds to counterclockwise rotation, whereas negative vorticity corresponds to clockwise rotation. To prevent negative and positive values from cancelling each other out within and/or between samples, the absolute value of vorticity was used at each point:

$$\left| \omega_{i,j} \right|=\left| \frac{V_{i+1,j}-V_{i-1,j}}{2\Delta x}-\frac{U_{i,j+1}-U_{i,j-1}}{2\Delta y} \right|$$

The average absolute value of all vorticity blocks (N) in a single frame was calculated as follows:

$$\overline{\left| \omega\right|}=\frac{1}{N}\sum_{i,j=1}^{N} \left| \omega_{i,j} \right|$$

Subsequently, this average was calculated for all frames (M number of frames and indexed by k):

$$\left\langle\left| \omega\right| \right\rangle=\frac{1}{MN}\sum_{k=1}^{M} \sum_{i,j=1}^{N} \left| \omega_{i,j}^{\left( k \right)} \right|$$

After measurements, medium containing the beads was aspirated and the monolayer was washed three times with PBS to remove residual beads. Subsequently, ciliated cell enrichment medium was added to the apical compartment.

**Apical-out nasal organoid rotation assay to measure 3D cilia activity (assay 3)**

Cilia-driven organoid rotation was measured in 3D AONOs generated from epithelial fragments of 2D differentiated HNECs. First, 2D differentiated monolayers were apically washed with pre-warmed ADMEM-F12 (Gibco) and basolaterally treated with collagenase type II solution diluted in ADMEM-F12 (1 mg/mL; Thermo Fischer Scientific, Cleveland, OH, USA). Cells were incubated for approximately 45 min at 37°C and 5% CO₂ until detachment of the epithelial sheet was observed. The dissociated epithelial layer was transferred to a 15 mL tube containing 1 mL ADMEM-F12 and mechanically fragmented by pipetting. Fragments were filtered through a 100 μm MACS® SmartStrainers cell strainer (Miltenyi Biotech B.V. & Co. KG, Bergisch Gladbach, Germany), centrifuged (206×*g*, 5 min, 4^o^C), and resuspended in 4 mL differentiation medium. Organoid formation proceeded for 1-4 days under continuous rotations in 15 mL filter cap tubes (CELLSTAR, Greiner Bio-One GmbH, Frickenhausen, Germany) on an orbital shaker at 37°C and 5% CO₂. After 1-4 days, organoids were sequentially filtered using 100 μm and 30 μm strainers. Organoids retained on the 30 μm filter were collected by inverting the filter into a new tube and washing with 10 mL ADMEM-F12 medium. Collected organoids were centrifuged (132×*g*, 5 min, 4°C) and embedded in 4 μL droplets of 0%, 25%, or 50% Matrigel (Corning) in pre-warmed 96-well tissue culture plates (Corning). Droplets were solidified for 20–30 min at 37°C and 5% CO₂ and 100 µL differentiation medium was added. Rotation of AONOs was recorded in bright-field using a 2.5x dry objective on a ZEISS Celldiscoverer 7 microscope (ZEISS) equipped with environmental control (37°C and 5% CO₂) for 30 s, acquiring 1 frame/s. Organoid segmentation was performed on the first frame using OrgaSegment, a deep-learning model originally developed for intestinal organoids [7]. Angular velocity of individual rotating organoids was quantified using an adapted motion-tracking algorithm [8] (<https://github.com/Living-Technologies/cilliary_function_analysis>).

Cilia-driven organoid rotation was validated using paclitaxel, a microtubule-stabilizing agent that interferes with ciliary motility [9, 10], at concentrations ranging from 6.25 μM to 200 μM applied for 6 h. Control organoids received equivalent concentrations of DMSO (Sigma-Aldrich). Organoid rotation was recorded before treatment (t = 0), immediately after treatment removal (t = 6 h), and 18 h after treatment removal (t = 24 h). Viability following treatment was assessed using CellTiter-Glo assay (Promega Corporation, Madison, USA) according to the manufacturer’s instructions, with luminescence as readout.

**RNA isolation, cDNA synthesis, and quantitative PCR**

Cells were collected, and total RNA was extracted from 2D differentiated HNECs using the NucleoSpin® RNA kit (VWR International, PA, USA) according to manufacturer’s instructions. Isolated RNA was stored at -80°C until further analysis. RNA concentration and purity were assessed using a DS-11 spectrophotometer (DeNovix Inc., Wilmington, USA), after which cDNA was synthesized using the iScript cDNA synthesis kit (Bio-Rad Laboratories, Inc., CA, USA) following the manufacturer’s protocol. Quantitative real-time PCR was carried out using specific primers for *TP63* (FW: CCACCTGGACGTATTCCACTG, REV: TCGAATCAAATGACTAGGAGGGG), *MUC5AC* (FW: ATTTTTTCCCCACTCCTGATG, REV: AAGACAACCCACTCCCAACC), and *FOXJ1* (FW: GGAGGGGACGTAAATCCCTA, REV: TTGGTCCCAGTAGTTCCAGC) with iQ SYBR Green Supermix (Bio-Rad Laboratories) on a CFX96 real-time detection system. Each reaction was initiated with a 3 min denaturation step at 95°C, followed by 40 cycles of 95°C for 10 s and primer-specific annealing at the appropriate temperature for 30 s. For relative gene expression analysis, Ct values were normalized to the housekeeping genes *ATP5B* (FW: TCACCCAGGCTGGTTCAGA, REV: AGTGGCCAGGGTAGGCTGAT), and *RPL13A* (FW: AAGGTGGTGGTCGTACGCTGTG, REV: CGGGAAGGGTTGGTGTTCATCC), selected for their stable expression in airway epithelial cells under varying experimental conditions. Relative expression levels were calculated using the 2-ΔΔCT method [11].

**Immunofluorescence staining of 2D differentiated cultures and 3D apical-out nasal organoids**

2D differentiated cultures on transwells and AONOs were fixed in 4% paraformaldehyde (Sigma-Aldrich) for 15 min, permeabilized with PBS containing 0.3% Triton X-100 (Sigma-Aldrich) for 30 min, and blocked in PBS with 3% BSA (Sigma-Aldrich) and 0.3% Triton X-100 for 60 min. Primary antibodies diluted in blocking buffer against β-tubulin IV (1:500, ab179509, Abcam, Cambridge, United Kingdom), MUC5AC (1:500, MA1-38223, Thermo Fisher Scientific), and/or DNAI1 (1:125, ab171964, Abcam) were applied for 1 h at room temperature (2D cultures) or overnight at 4°C (AONOs). After three washes with PBS, samples were incubated with secondary antibodies diluted in blocking buffer, including Alexa Fluor 488 (1:500; A-11034, Thermo Fisher Scientific), Alexa Fluor 647 (1:500; A-21240, Thermo Fisher Scientific), Phalloidin (1:250, A-34055,Thermo Fisher Scientific), and/or DAPI (1:1000, D9542, Sigma) for 30 min at room temperature in the dark (2D cultures) or overnight at 4°C (AONOs). Samples were subsequently washed three times with PBS. AONOs were stored in PBS at 4°C until imaging. Transwell membranes were excised and mounted on microscope slides using ProLong Gold antifade reagent without DAPI (Thermo Fischer Scientific). IF imaging was performed using a Leica THUNDER imager (Leica Microsystems, Vienna, Austria) at 40x magnification or a Zeiss LSM800 confocal microscope (ZEISS, Oberkochen, Germany) at 63x magnification.

**Basolateral LNP-mRNA delivery**

HNECs from healthy controls or individuals with PCD harboring a *DNAI1* variant were differentiated on 0.4 μm and 1.0 μm transwell inserts (Falcon® Permeable Support for 24-well Plate with 1.0 µm Transparent PET Membrane, Falcon, Göteborg, Sweden) as described above. After 14 days of differentiation, monolayers were treated basolaterally with LNPs containing *tdTomato* mRNA (5 or 10 μg/mL, ReCode Therapeutics). Based on the resulting transfection efficiency, 1.0 μm pore size transwells were used for all subsequent LNP-based assays. From day 14 onward, cultures were treated three times per week with LNPs containing *DNAI1* mRNA (ReCode Theapeutics) at concentrations of 0 µg/mL, 2.5 µg/mL, 5 µg/mL, or 10 µg/mL. LNPs were diluted in differentiation medium and added to the basolateral compartment. After 5 h, the LNP-containing medium was aspirated, the basolateral compartment was washed with PBS, and fresh differentiation medium supplemented with DAPT and DMH-1 was added.

**Western blot**

For Western blot analysis of DNAI1, differentiated HNECs were lysed in RIPA Buffer (Thermo Fisher Scientific). Lysates were homogenized using QIAshredder columns (Qiagen N.V., Hilden, Germany) and centrifuged at 20,000×*g* for 5 min. Protein concentration was determined using the Pierce BCA Protein Assay Kit (Thermo Fisher Scientific), according to manufacturer’s instructions. SDS-PAGE was performed using 30 µg protein per lane, followed by transfer onto polyvinylidene difluoride membranes (Immobilon FL, MO, USA) overnight at 30 V and 4^o^C. Membranes were blocked with 5 % (w/v) non-fat milk powder (Campina, Amersfoort, Netherlands) dissolved in TBS-T for 1 h at room temperature. Membranes were then incubated with primary antibodies against DNAI1 (1:2000, ab171964, Abcam) and β-tubulin IV (1:7500, ab179509, Abcam, loading control) diluted in 0.5 % (w/v) non-fat milk powder in TBS-T for 3 h at room temperature. Proteins were detected using a swine anti-rabbit-HRP-conjugated secondary antibody (1:2000, P0217, Dako, Glostrup, Denmark) diluted in 0.5% (w/v) non-fat milk powder/TBS-T and incubated for 1 h at room temperature. Chemiluminescent signals were acquired using a Bio-Rad ChemiDoc Touch Imaging System (version 3.0.1; Bio-Rad Laboratories).

**Statistical analysis**

All experiments were performed using a subset of HNECs derived from 8 healthy individuals and 13 individuals with PCD. Experiments were performed in technical duplicates or triplicates. Data are presented from three biological replicates, either from a single individual or pooled from different individuals having a similar background (e.g. healthy controls). Data are presented as mean ± SEM. Z-scores were calculated using the following formula: z = (x - μHC)/SDHC, where x is the measured value and μHC and SDHC are the mean and SD of the healthy controls, respectively. Statistical analyses were performed using Graphpad Prism 10.4.1 software and R Studio. Depending on the dataset, significance was assessed using appropriate parametric or non-parametric tests, as indicated in the figures’ legends. Statistical significance was defined as * p < 0.05, ** p < 0.01, *** p < 0.001, and **** p < 0.0001.

**Data availability**

The raw datasets generated and/or analyzed during the current study are available from the corresponding author upon reasonable request.

**Code availability**

The BT assay analysis code and the adapted motion-tracking algorithm code [8] to analyze AONO rotation are available at GitHub: <https://github.com/Living-Technologies/cilliary_function_analysis>

**Supplementary Table 1.** Overview of clinical diagnostics of the biallelic PCD genetic alterations described in the current study

| PCD  codename | Gene | cDNA Change (Protein Change) | Nasal NO  (nl/min) | HSVM before cell culture | TEM defect |
| --- | --- | --- | --- | --- | --- |
| PCD01 | *CCDC39* | c.357+1G>C (unknown) / c.2347_2351del  (p.F783Yfs*3) | 4 | Immotile cilia,  low number of cilia | Microtubular disorganization and IDA defect |
| PCD02 | *CCDC39* | homozygous c.2158+1G>A  (unknown) | N/A | Normal to increased frequency, stiff moving ciliary tip, varying coordination | Microtubular disorganization and IDA defect |
| PCD03 | *CCDC40* | c.248del  (p.A83Vfs*84) | N/A | N/A | N/A |
| PCD04 | *CCDC40* | c.248del  (p.A83Vfs*84) /  1855C > T  (p.Q619X) | 4 | Low frequency, low amplitude and no coordination | Microtubular disorganization and IDA defect |
| PCD05 | *CCNO* | c.793dup (p.V265Gfs*106) | N/A | Ciliary agenesis | N/A |
| PCD06 | *DNAH5* | c.[10384C > T]  (p.Q3462X) /  [13338+5G > A]  (unknown) | N/A | Immotile cilia | ODA/IDA defect |
| PCD07 | *DNAH5* | p.T2966M, p.V4313M  c.7408-2A>G. | N/A | N/A | N/A |
| PCD08 | *DNAH11* | c.2712G>A (unknown)/  unknown | 3 | Mostly immotile cilia, some with increased frequency and low amplitude and no coordination | Normal |
| PCD09 | *DNAH11* | c.9824A>C  (p.Y3275S) /  c.13304-1G>A  (unknown) | 18 | Normal frequency, low amplitude, varying coordination | Normal |
| PCD10 | *DNAI1* | c.48+2dup  (p.S17Vfs*9) /  c.1640 [T>C]  (p.V547A) | 17 | N/A | N/A |
| PCD11 | *DNAI1* | c.1431del (p.K477Nfs*2) | 2 | N/A | N/A |
| PCD12 | *DNAI1* | c.48+2dup  (p.S17Vfs*9) | N/A | Immotile cilia and cilia with low frequency, low amplitude and no coordination | ODA/IDA defect |
| PCD13 | *HYDIN* | c.8356[C>T] (p.R2786X) / c.13867[G>T] (p.G4623X) | N/A | Low frequency, low amplitude, no coordination | Normal |

HSVM: high speed video microscopy; IDA: inner dynein arm; N/A: not applicable; NO: nasal nitric oxide; ODA: outer dynein arm; PCD: Primary Ciliary Dyskinesia; TEM: transmission electron microscopy

**Supplementary Table 2.** Components of isolation and expansion medium

| Basic expansion medium | | |
| --- | --- | --- |
| Compound | **Concentration** | **Identifier, Source** |
| BEpiCM-b | 50% (v/v) | Cat#3211, Sciencell |
| ADMEM/F-12 | 44% (v/v) | Cat#12634-028, Thermo Fisher Scientific |
| B-27 Supplement, serum free | 2% (v/v) | Cat#17504001, Thermo Fisher Scientific |
| HEPES | 10 mM | Cat#15630080, Thermo Fisher Scientific |
| GlutaMAX supplement | 1% (v/v) | Cat#35050-061, Thermo Fisher Scientific |
| Penicillin/streptomycin | 1% (v/v) | Cat#15070-063, Thermo Fisher Scientific |
| Hydrocortisone | 0,5 µg/mL | Cat#H0888, Sigma-Aldrich |
| N-Acetyl-L-cysteine | 1,25 mM | Cat#A9165, Sigma-Aldrich |
| Primocin | 100 µg/mL | Cat#ant-pm-2, InvivoGen |
| A83-01 | 1 µM | Cat#2939/10, Tocris |
| (±)- Epinephrine hydrochloride | 0,5 µg/mL | Cat#E4642, Sigma-Aldrich |
| Y-27632 | 5 µM | Cat#S1049, Selleck Chemicals |
| RSPO3-Fc Fusion Protein conditioned medium | 2% | Cat#R001 - 100 mL, U-Protein Express |
| Human Heregulin-beta 1 | 5 nM | Cat#100-03, PeproTech |
| rhFGF10 | 100 ng/mL | Cat#100-26, PeproTech |
| rhHGF | 25 ng/mL | Cat#100-39H, PeproTech |
| Isolation Medium | | |
| Basic expansion medium | 100% (v/v) |  |
| Amphotericin B | 250 µg/mL | Cat#15290018, Thermo Fisher Scientific |
| Gentamicin | 50 µg/mL | Cat#G1397, Sigma-Aldrich |
| Vancomycin | 50 µg/mL | Cat#SBR00001, Sigma-Aldrich |
| Expansion Medium | | |
| Basic expansion medium | 100% (v/v) |  |
| DAPT | 5 µM | Cat#15467109, Thermo Fisher Scientific |
| Rapamycin | 5 µM | Cat#553210-1MG, Sigma |

ADMEM-F12: Advanced Dulbecco's Modified Eagle Medium/Nutrient Mixture F-12; BEpiCM: Bronchial Epithelial Cell Medium; rhFGF10: recombinant human Fibroblast growth factor 10; rhHGF: recombinant human Hepatocyte growth factor

**Supplementary Table 3.** Components of differentiation medium

| Differentiation medium | | |
| --- | --- | --- |
| Compound | Concentration | Identifier, Source |
| ADMEM-F12 | 98,5 % (v/v) | Cat#12634-028, Thermo Fisher Scientific |
| 3,3′,5-Triiodo-L-thyronine sodium salt | 100 nM | Cat#T6397, Sigma-Aldrich |
| Hydrocortisone | 0.5 µg/ml | Cat#H0888, Sigma-Aldrich |
| (±)-Epinephrine hydrochloride | 0.5 µg/ml | Cat#E4642, Sigma-Aldrich |
| A83-01 | 50 nM | Cat#2939/10, Tocris |
| TTNPB | 100 nM | Cat#16144-1, Cayman |
| rhEGF | 0.5 ng/mL | Cat#AF-100-15, PeproTech |
| Penicillin/streptomycin | 1% (v/v) | Cat#15070-063, Thermo Fisher Scientific |
|  | **Ciliated cell enrichment medium** | |
| Differentiation medium | 100 % (v/v) |  |
| DAPT | 5 μM | 15467109, Thermo Fisher Scientific |
| DMH-1 | 500 nM | S7146, Selleck Chemicals |

ADMEM-F12: Advanced Dulbecco's Modified Eagle Medium/Nutrient Mixture F-12; DMH-1: dorsomorphin homolog 1; rhEGF: recombinant human Epidermal Growth Factor

**Supplementary Figure Legends**

**Supplementary Figure 1 | Cellular and mRNA composition of the pseudostratified epithelium changes during cilia-promoting low liquid-liquid interface differentiation.**

**A** Graphical illustration of the experimental timeline showing six endpoint measurements during low liquid-liquid interface differentiation with DAPT and DMH-1. **B** Representative IF images of low liquid-liquid interface cultures (medium containing DAPT and DMH-1) derived from a healthy control, stained for the ciliated cell marker β-tubulin IV (red), and the secretory cell marker MUC5AC (cyan) after days 0, 7, 14, 21, 28, and 42 of differentiation. **C** Quantification of β-tubulin IV and **D** MUC5AC signal. **E** Quantitative PCR comparing the mRNA expression of *MUC5AC,* **F** *FOXJ1* and **G** *TP63* after 0, 7, 14, 21, 28, and 42 days of differentiation. Results are shown as mean ± SEM of HNECs from three healthy individuals. Scale bars, 50 µm. * p < 0.05 and **** p < 0.0001 as compared to differentiation day 0 in panels **C, E, G** (Kruskal-Wallis test with Dunn’s multiple comparisons), and to differentiation day 0 in panels **D, F** (one-way ANOVA with Dunnett’s multiple comparisons).

**Supplementary Figure 2 | Ciliary beat frequency of healthy differentiated HNECs is independent of ciliary active area.**

Correlation graph between active ciliated area and ciliary beat frequency from data shown in **Figure 2 C, D** (r*^2^* = 0.1062). A Spearman correlation test was performed.

**Supplementary Figure 3 | Optimization and validation of AONO rotation.**

**A** Representative IF images of AONOs derived from PCD (PCD01, PCD10) individuals. Nuclear markers are stained with DAPI (gray), and ciliated-cell markers with β-tubulin IV (red). Scale bar, 25 μm. **B** Segmentation of individual organoids derived from a healthy control seeded using various Matrigel concentrations (0 %, 25 % and 50 %) and seeding timepoints (24, 48, 72, and 96 h). Scale bar, 50 μm. **C** Absolute number, **D** percentage and **E** angular speed of rotating organoids using various Matrigel concentrations (0 %, 25 % and 50 %) and seeding timepoints (24, 48, 72, and 96 h). Angular speed was measured in rad/s. **F** Cell viability of AONOs (% of control) after paclitaxel treatment measured using the CellTiter-Glo assay. Results are shown as mean ± SEM of HNECs from one healthy individual. * p < 0.05, and **** p < 0.0001 as compared to 24 h for each Matrigel concentration in panel **D** (two-way ANOVA) and **E** (Welch-type two-way ANOVA), and in panel **F** a linear regression curve was fitted (nested-model ANOVA).

**Supplementary Figure 4 | Assay-dependent differences in ciliary activity distinguish healthy and PCD individuals.**

**A** Quantification of the percentage of active ciliated area (%), **B** CBF (Hz) within the active areas, **C** bead velocity (μm/s) and **D** vorticity (s^-1^) in low liquid-liquid interface cultures from five healthy and 11 PCD individuals. **E** Quantification of rotating AONOs (%) in the Matrigel droplets and **F** angular speed (rad/s) of rotating organoids from five healthy and 11 PCD individuals. Three biological replicates (three consecutive passages of HNECs) were analyzed. Results are shown as mean ± SEM of HNECs from five healthy and 11 PCD individuals. * p < 0.05 and ** p < 0.01 as compared to healthy controls in panels **A-F** (Mann-Whitney U-test without Holm’s correction, significance of non-adjusted p values is shown).

**Supplementary Figure 5 | Z-score normalization and combined assay analysis reveal limited intra-PCD differences but clear separation from healthy controls.**

**A** Z-score of each indicated cilia functional measurement for the 11 PCD-derived HNECs normalized to healthy controls. **B** Box plots show averaged z-score for 2D cilia activity (assay 1), **C** 2D bead transport (assay 2), and **D** 3D cilia activity (assay 3) for the 11 PCD-derived HNECs normalized to healthy controls. **E** Correlation graph between 2D bead transport (assay 2) and 3D cilia activity (assay 3) shown as z-score values for the seven different tested PCD genes. Results are shown as individual points in boxplots of HNECs from five healthy and 11 PCD individuals. * p < 0.05 and *** p < 0.001 compared to healthy controls in panels **B-D** (Mann-Whitney U-test).

**Supplementary Figure 6 | Delivery efficiency of single *TdTomato* mRNA-LNP treatment and its effect on ciliary activity.**

**A** Graphical illustration and **B** representative IF images showing red fluorescence after single basolateral *tdTomato* mRNA-LNP treatment on both healthy and PCD-derived differentiated HNECs (PCD09) on 0.4 or 1 μm pore size filters. **C** Active ciliated area (%) and **D** CBF (Hz) within the active areas of low liquid-liquid interface differentiated healthy (n = 2) and PCD-derived (PCD09) HNECs upon single basolateral treatment with different *tdTomato* mRNA-LNP concentrations. One biological replicate was analyzed. Scale bar, 50 μm

**Supplementary Figure 7 | Outcome parameters of 2D cilia activity and 2D bead transport show correlation in *DNAI1* mRNA LNP-treated HNECs harboring a *DNAI1* alteration.**

**A** Correlation graph between bead velocity (μm/s) and vorticity (s^-1^)  (r*^2^* = 0.8924), **B** active ciliated area (%) and bead velocity (μm/s) (r*^2^* = 0.6921) and **C** active ciliated area (%) and bead vorticity (s^-1^) (r*^2^* = 0.7574) from data shown in **Figure 7B, E, F**. A Spearman correlation test was performed for panels **A-C**.
