## Supplementary Figure for "Personalized multi-assay profiling of respiratory motile ciliopathies and mRNA therapy"

Supplementary Figure 1

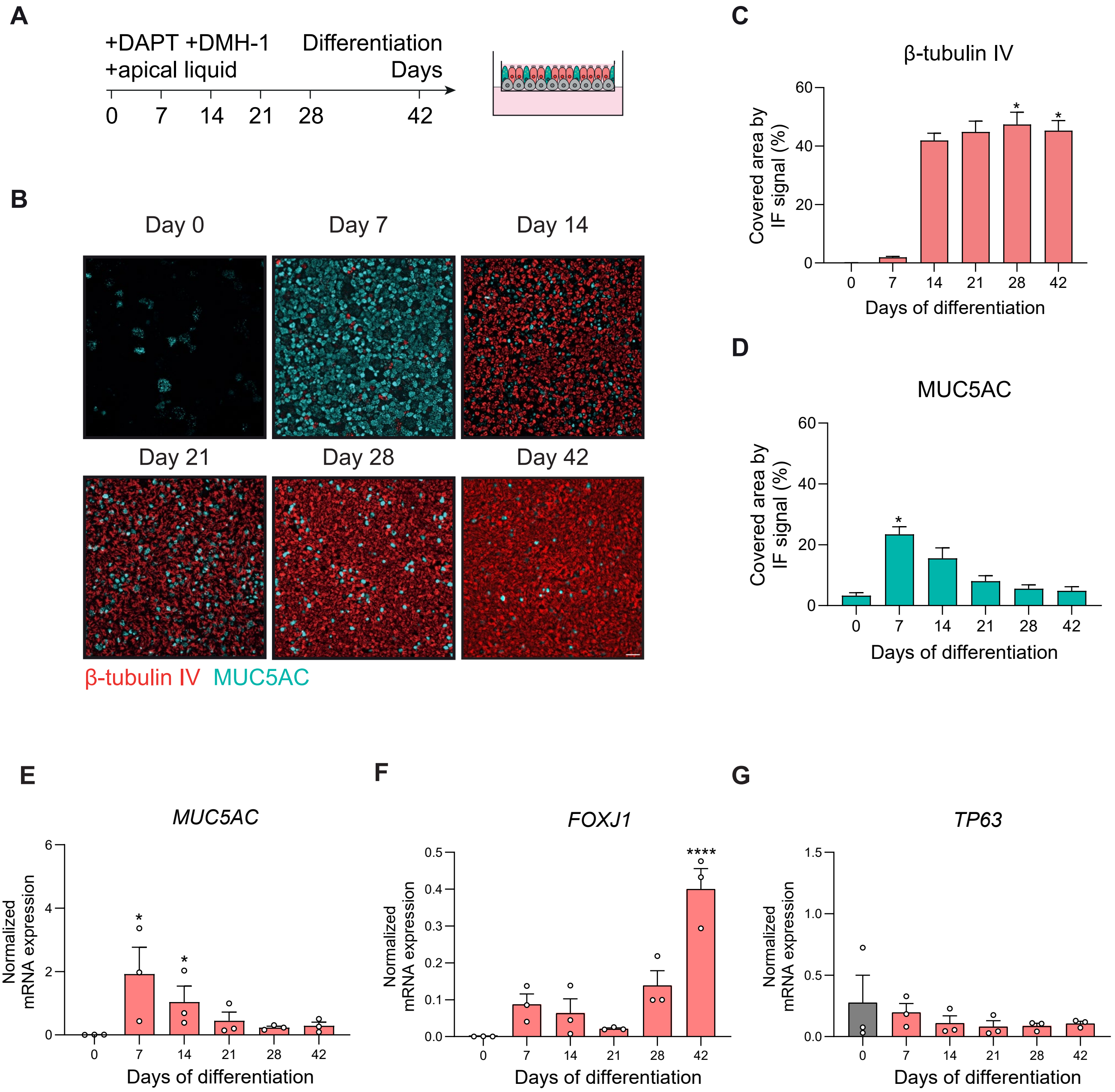

### Supplementary Figure 2

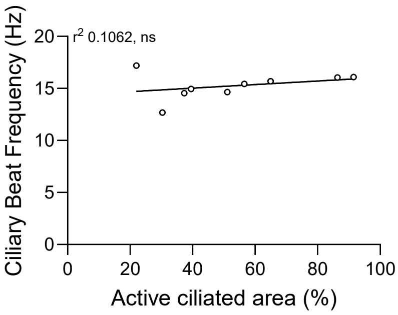

Supplementary Figure 3

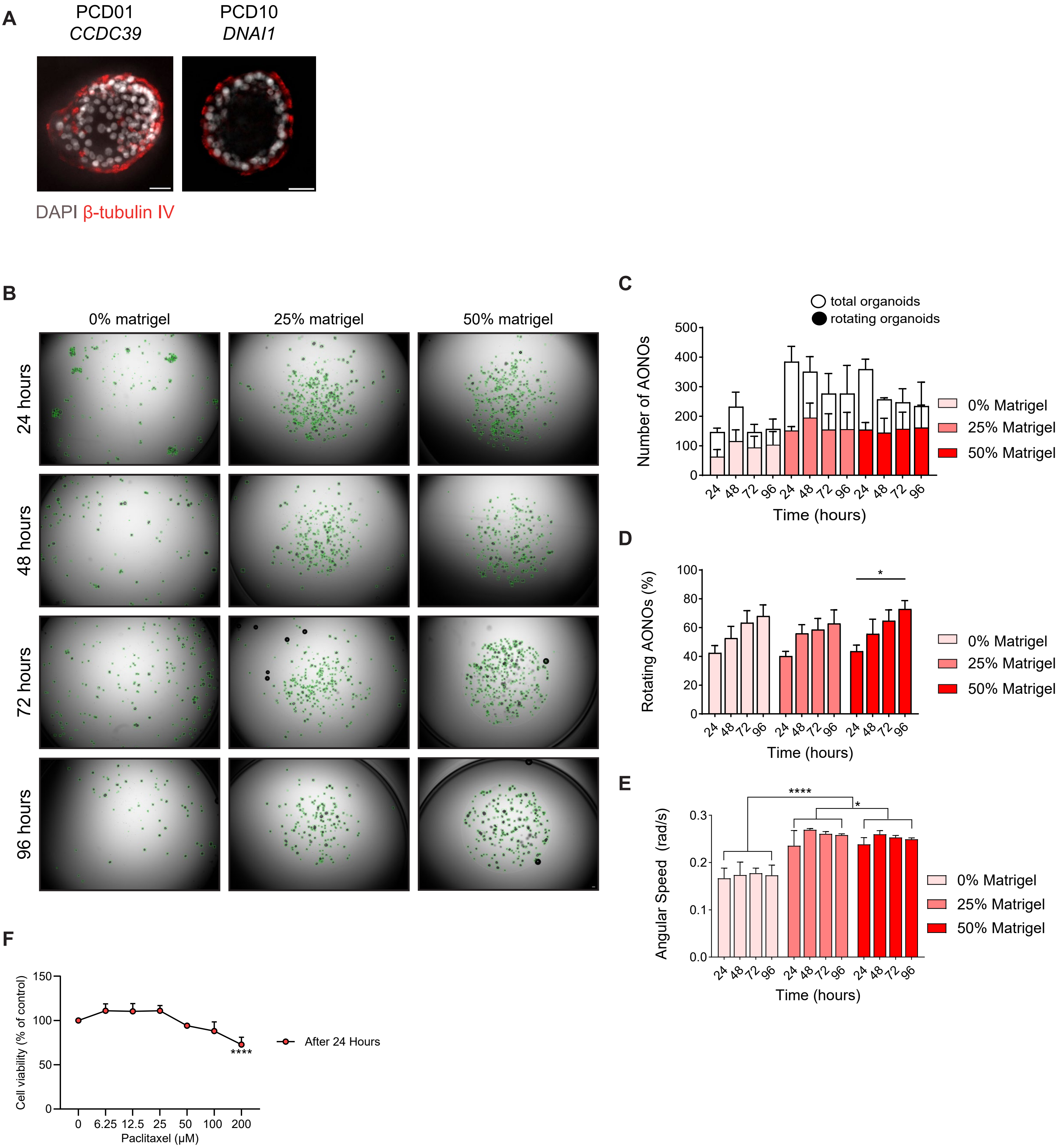

Supplementary Figure 4

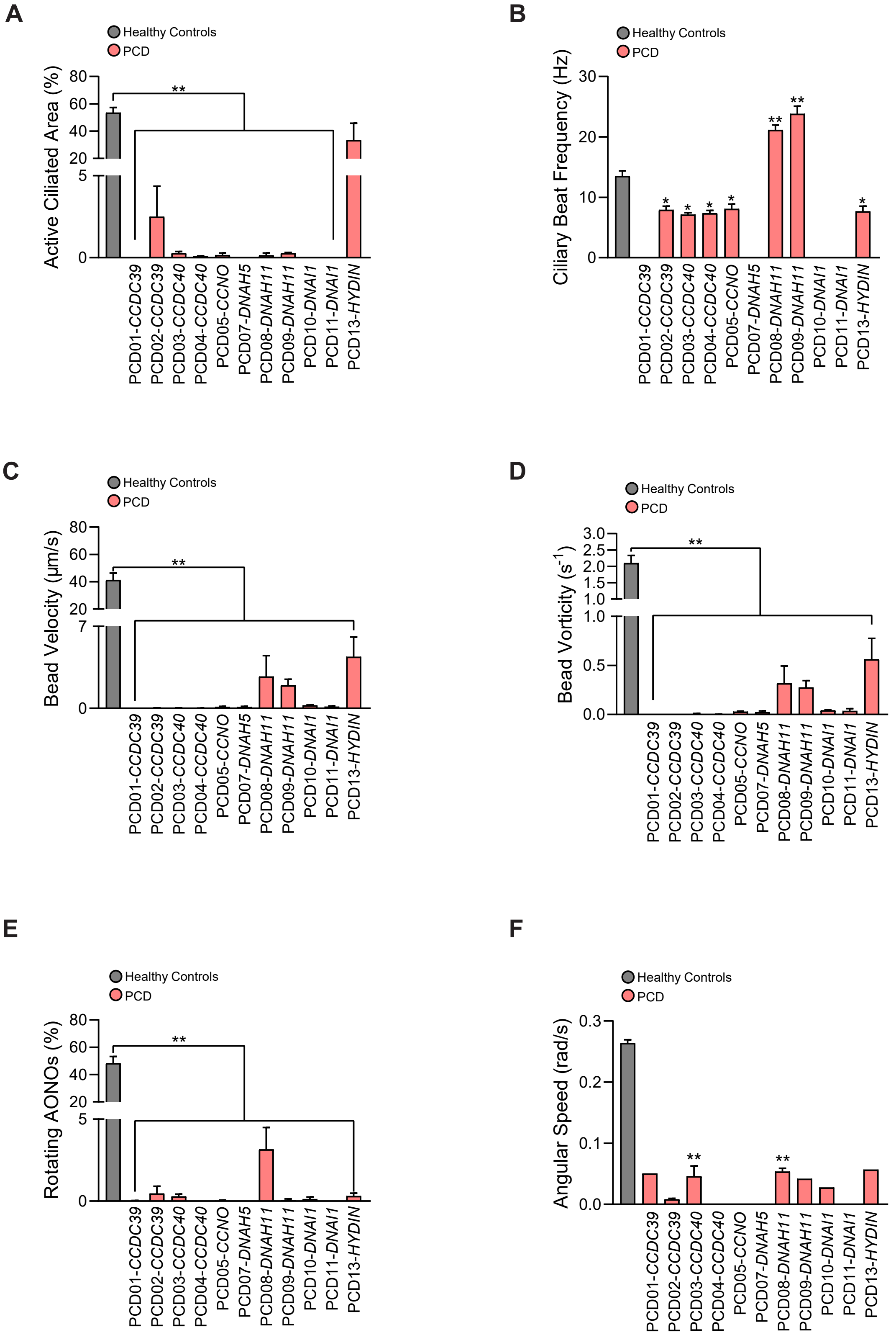

Supplementary Figure 5

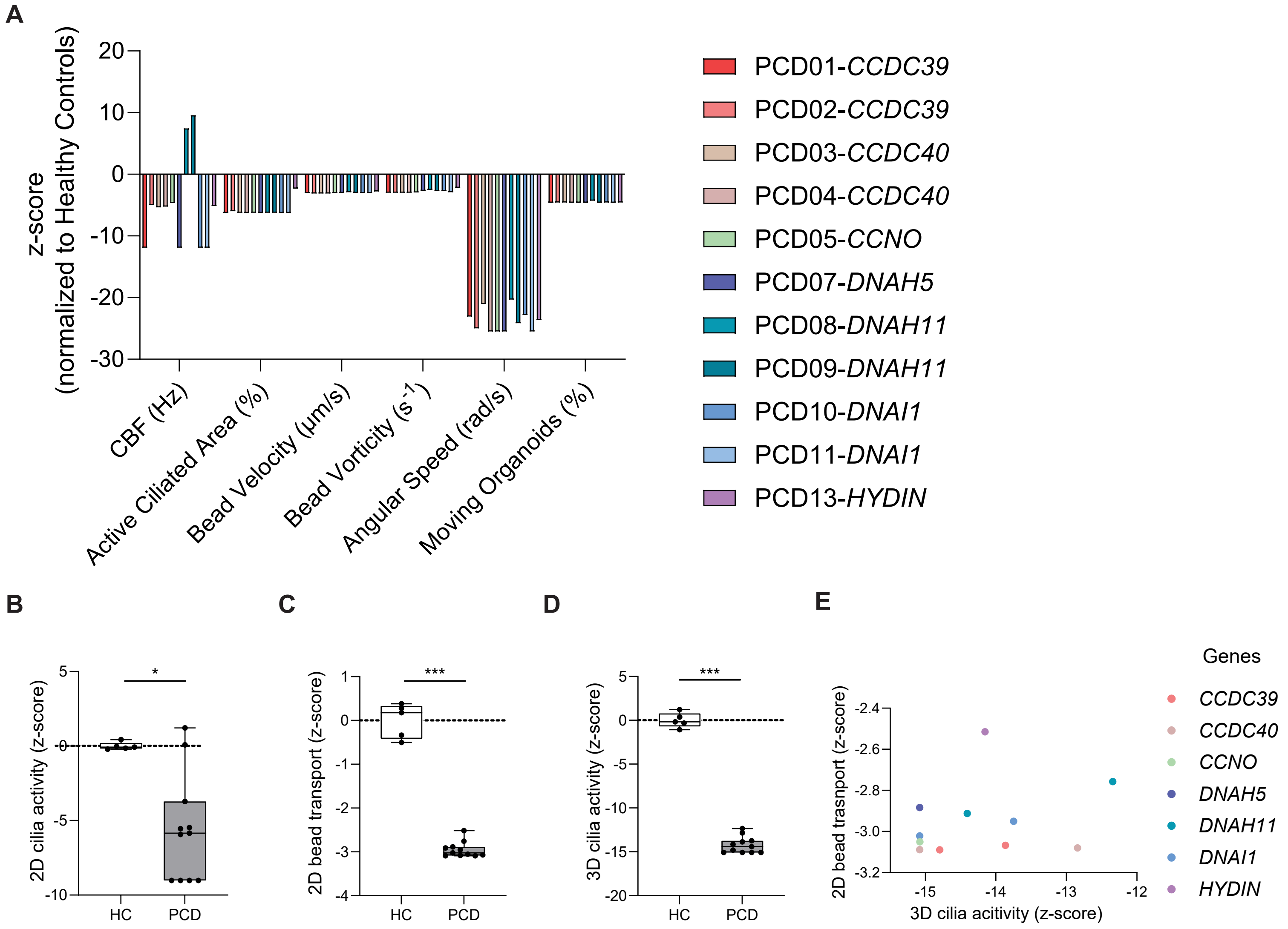

Supplementary Figure 6

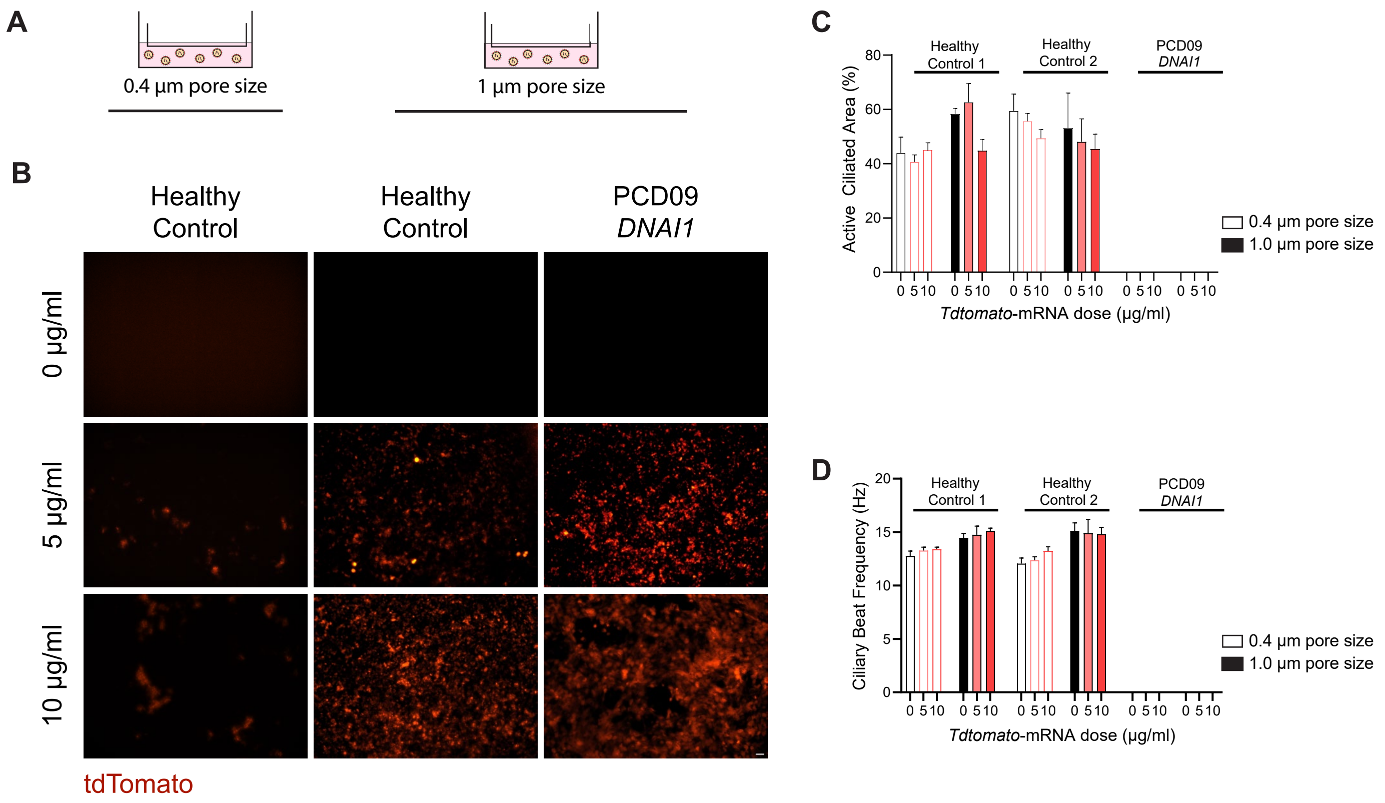

Supplementary Figure 7

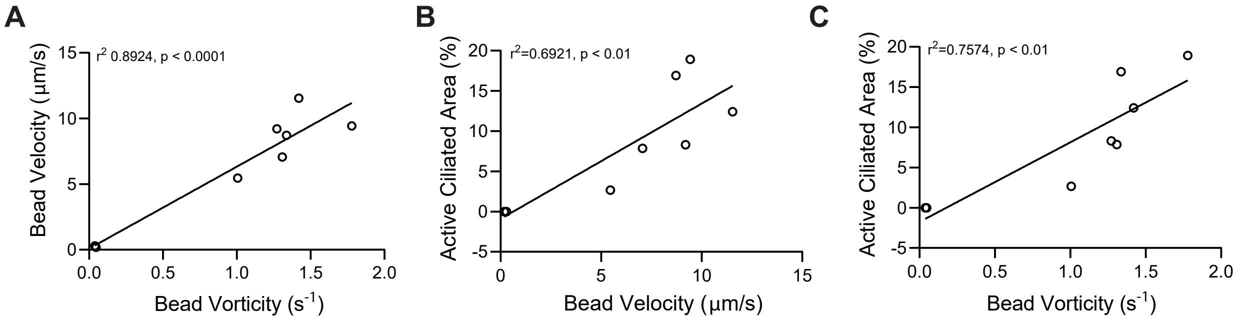
